# Siderophore-Mediated Competition Between Cystic Fibrosis Pathogens Increases Virulence Factor Production

**DOI:** 10.64898/2026.10.23.761465

**Authors:** Niladri Bhowmik, Laura Rodriguez-Velandia, Mercedes Pérez Pérez, Sydney M. Morabbi, Laura M. Sanchez, Reed M. Stubbendieck

## Abstract

The airways of people with cystic fibrosis (CF) are colonized by a polymicrobial community and coinfections with multiple pathogens may exacerbate disease. By analyzing 16S rRNA gene sequencing datasets, we determined that *Achromobacter*-*Pseudomonas* coinfection was associated with worse lung function than infection with either pathogen alone, or coinfections with *Pseudomonas* and other CF bacterial pathogens. We discovered *Achromobacter* spp. induced production of the siderophore pyoverdine by *Pseudomonas aeruginosa* strain PAO1 and other clinical isolates on synthetic CF sputum medium (SCFM), supplemented with physiologically relevant iron concentrations. We then tested 21 *Achromobacter* strains, encompassing seven species, and determined that 17 (81%) induced pyoverdine production. We sequenced the full genomes of these *Achromobacter* and found 18 (86%) encoded a biosynthetic gene cluster for the production of alcaligin. This compound is an iron-chelating siderophore that has previously been shown to be involved in colonization of the respiratory tract by *Bordetella* spp. Using mass spectrometry imaging, we confirmed that *Achromobacter dolens* strain RMS0405 produces alcaligin during coculture with PAO1. Further, we found that supplementing SCFM with increasing iron abolishes pyoverdine induction by PAO1 during coculture. Pyoverdine is known to contribute directly to virulence and regulate expression of other virulence factors. Using transcriptomics and proteomics, we determined that RMS0405 induces a low-iron stress response in PAO1 and stimulates the production of exotoxin A and other known virulence factors. Together, our results indicate that *Achromobacter* spp. and *P. aeruginosa* may exacerbate virulence in the airways of people with CF as a byproduct of competition for iron.

**Importance:** Within polymicrobial infections, competition between pathogens is thought to attenuate virulence, as microbes direct resources away from pathogenesis and toward microbial interactions. However, by coculturing the cystic fibrosis pathogens *Achromobacter* spp. and *Pseudomonas aeruginosa*, we determined that competition induces pyoverdine production by the latter pathogen. This secondary metabolite functions in iron acquisition, is cytotoxic toward eukaryotic cells, and activates production of other virulence factors. We demonstrated that production of the siderophore alcaligin by *Achromobacter dolens* imposes iron limitation and induces pyoverdine production by *P. aeruginosa*. Our finding challenges a conventional view that competition limits pathogenic potential, and instead suggests that interactions between coinfecting microbes can amplify virulence. Notably, disrupting the signals or pathways that drive competitive upregulation of virulence factors could attenuate disease severity without relying on traditional antibiotics. This approach could open new avenues for understanding pathogenesis and for developing virulence factor-targeting therapeutics tailored to polymicrobial infections.

## Introduction

Cystic fibrosis (CF) is a multisystem disease caused by mutations in the gene encoding the CF transmembrane conductance regulator (*CFTR*). These mutations disrupt chloride ion transport activity, resulting in the accumulation of a thick and adherent mucus that provides a niche for microbial colonization (1, 2). Impaired mucociliary clearance promotes chronic infection, during which pathogens in the airways of people with CF (pwCF) produce virulence factors that damage epithelial tissues, leading to lung function decline, and, ultimately, death. Despite recent advancements in treatment, including development of CFTR modulators, such as elexacaftor/tezacaftor/ivacaftor (ETI), CF remains a life-shortening disease with significant morbidity and mortality, reflected in a median age of death of 37 years (3).

The most well-characterized pathogen in the airways of pwCF is *Pseudomonas aeruginosa*. Advances in metagenomics and culturomics revealed that the airways of pwCF are colonized by a complex microbiota that includes other oropharyngeal and pathogenic bacteria (4–6). We recently showed that these two groups of bacteria colonize the CF airway with distinct patterns. Oropharyngeal genera are prevalent but rarely abundant, whereas pathogens are less prevalent but often reach high relative abundance when present. These microbiota states are associated with different clinical outcomes for pwCF (7). Our data further suggested that pathogens come to dominate in part by producing bioactive metabolites to inhibit other members of the CF airway microbiota (7). Among CF pathogens, the roles that bioactive metabolites play in interactions between *P. aeruginosa* and *Staphylococcus aureus* have been well established (8–11). However, the consequences of these metabolite-mediated interactions between *P. aeruginosa* and other CF pathogens, and their impact on disease, are less understood.

*Achromobacter* spp. are Gram-negative bacteria commonly found within both terrestrial and aquatic environments. These bacteria have emerged as pathogens in the airways of pwCF over the last 30 years, rising globally in incidence from approximately 2-3% in the 1990s to approximately 7-10% in the early 2020s (12). The type species is *Achromobacter xylosoxidans*, however early studies did not distinguish between this and ten other *Achromobacter* spp. that are now known to infect the airways of pwCF (13–15). Infections with *Achromobacter* are associated with a strong inflammatory response due to the production of siderophores, type III secretion, and type VI secretion systems (16–23).

*Achromobacter* infections are associated with severe loss-of-function *CFTR* genotypes, including those that do not respond to ETI, and are difficult to treat due to the multidrug resistance of these pathogens. Thus, infection with *Achromobacter* remains a threat to pwCF with the fewest therapeutic options (23–26).

Approximately half of pwCF with chronic *Achromobacter* infection are coinfected with *P. aeruginosa.* These coinfections are frequently associated with more severe clinical outcomes and greater annual decrease in pulmonary health (16–19, 23, 27). Few studies have directly characterized interactions between *Achromobacter* spp. and *P. aeruginosa*. For instance, Menetrey et al. tested 53 *A. xylosoxidans*-*P. aeruginosa* pairs in coculture assays, finding eleven instances of growth inhibition, two instances of motility reduction, and seven instances of pigment reduction in *P. aeruginosa* (28). Besse and Menetrey et al. further showed that *A. xylosoxidans* strain Ax 200 altered the *P. aeruginosa* proteome, reducing production of proteins involved in swimming, biofilm formation, and siderophore production. Accordingly, the mortality of *Danio rerio* embryos coinfected with *A. xylosoxidans* Ax 200 and *P. aeruginosa* was halved, compared to infection with *P. aeruginosa* alone (29). Similarly, Sahl et al. recently reported that early-infection CF isolates of *A. xylosoxidans* inhibit *P. aeruginosa* biofilm formation *in vitro* (30). Although the mechanisms underlying virulence attenuation and biofilm inhibition remain unclear, Soh et al. previously demonstrated that 12 of 17 CF-associated *A. xylosoxidans* strains they tested inactivated the *Pseudomonas* quinolone signal (31), which is necessary for these virulence phenotypes (32). In addition to antagonism, Menetrey et al. also identified nine instances of growth promotion, eight instances of motility increase, and one instance of increased pigment production by *P. aeruginosa* during coculture with *A. xylosoxidans* (28). Thus, *Achromobacter* spp. can alter virulence-associated phenotypes in *P. aeruginosa* positively and negatively, but the specific mechanisms underlying these interactions are not well-understood.

In this study, we show that *Achromobacter*-*Pseudomonas* coinfection is associated with more severe disease than infection with either pathogen alone. During coculture, *Achromobacter* spp. impose iron limitation that induces production of pyoverdine, activates an iron-starvation response, and increases production of extracellular virulence factors by *P. aeruginosa*. Together, these data suggest that CF pathogens engage in exploitation competition for iron in the airways, which may lead to collateral damage to the host.

## Results

### Coinfections with *Achromobacter* and *Pseudomonas* exacerbate lung function outcomes in pwCF

We wanted to assess the impact of coinfection with *Achromobacter* and *Pseudomonas* on lung function through a reanalysis of 16S rRNA gene amplicon sequencing studies (7) (**Table S1** in the supplemental material). We limited our analyses to the genus level, because the 16S rRNA gene possesses variable and low resolution below this taxonomic level (33, 34).

To evaluate lung function, we used the forced expiratory volume in 1 s percent predicted (FEV_1_ pp) metric. We grouped samples based on whether they contained no pathogens (n=49), *Achromobacter* alone (n=43), *Pseudomonas* alone (n=709), or both pathogens together (n=58), using a 1% relative abundance threshold. There were significant differences in lung function across the groups (Kruskal-Wallis χ^2^ = 16.3, df = 3, *P* < 0.001). Relative to when no pathogen was detected, pwCF with either *Achromobacter* or *Pseudomonas* alone had FEV_1_ pp values about 9 to 10 points lower, whereas those with both pathogens had values lower by about 20 points (**Fig. 1A**). Across relative abundance thresholds ranging from 0.1% to 5%, this relationship held (**Fig. S1A** in the supplemental material). We found that higher combined abundance of the two pathogens was associated with lower FEV_1_ pp values. Further, pwCF whose airway microbiota was dominated (combined relative abundance >50%) by *Achromobacter* and *Pseudomonas* had the lowest FEV_1_ pp values (median 42.0 FEV_1_ pp, interquartile range [IQR]: 27 to 52 FEV_1_ pp), compared to those in which the two pathogens made up a smaller fraction (<50% relative abundance) of the microbiota (median: 68.5 FEV_1_ pp, IQR: 45 to 86 FEV_1_ pp) (Wilcoxon rank-sum test *P* = 0.018) (see **Fig. S1B** in the supplemental material).

**Figure 1.**
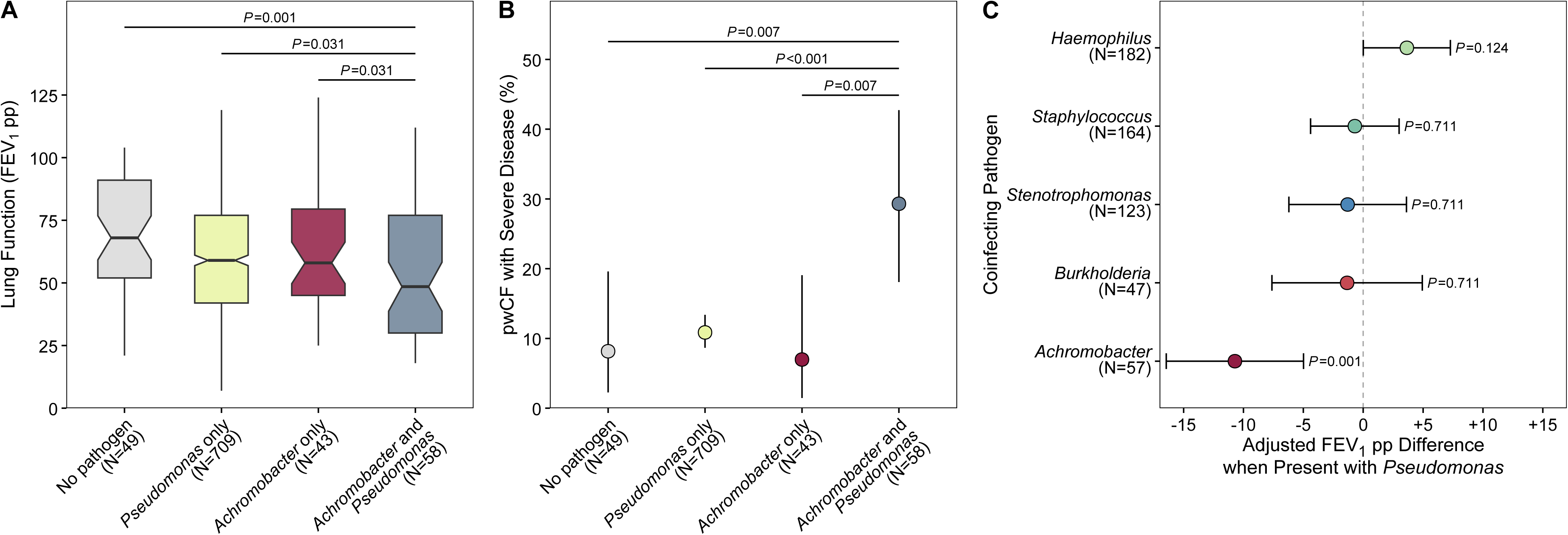
pwCF coinfected with *Achromobacter* and *Pseudomonas* exhibit more severe lung function outcomes. (**A**) Notched boxplot comparing lung function (FEV_1_ pp) between pwCF without any detected pathogens (<1% relative abundance) and pwCF grouped based on pathogen presence (defined as ≥1% relative abundance). The upper and lower bounds of the notched box plots indicate the 75th and 25th percentiles, respectively. The horizontal black bars and notches represent the median and 95% confidence interval (CI) of the median, respectively. The whiskers extend from the bounds of the box to the largest and smallest values that are no further than 1.5× the IQR. Horizontal lines indicate pairwise comparisons against the both-pathogens group, with Benjamini-Hochberg (BH)-adjusted *P*-values from Wilcoxon rank-sum tests. (**B**) Proportion of pwCF in each group with severe disease (FEV_1_ pp ≤ 30%). Points and lines show the proportion and 95% binomial CI. Horizontal lines indicate pairwise comparisons against the both-pathogens group, with BH-adjusted *P*-values from Fisher’s exact test. (**C**) Association of coinfection of *Pseudomonas* and other CF pathogens on lung function after adjustment for age and BioProject. Samples missing either of these values were excluded. Points and error bars show the average change in adjusted lung function and 95% CI. Each estimate is the change relative to *Pseudomonas*-positive samples lacking the corresponding pathogen, with the other pathogens held constant. The dashed vertical line marks no difference. BH-corrected *P*-values from per-coefficient *t*-tests are indicated to the right of each comparison.

The proportion of pwCF with severe disease differed between the four groups (χ^2^ = 19.3, df = 3, *P* < 0.001). Specifically, we found that pwCF coinfected with both pathogens were three to fourfold more likely to have severe lung disease (defined as FEV_1_ pp ≤ 30%) than those with *Achromobacter* or *Pseudomonas* alone (**Fig. 1B**). This effect was specific to *Achromobacter*, as coinfection with *Burkholderia*, *Haemophilus*, *Staphylococcus*, or *Stenotrophomonas* did not significantly alter lung function, relative to *Pseudomonas* alone (**Fig. 1C**). Thus, these data support clinical studies concluding that coinfection with *Achromobacter* spp. and *P. aeruginosa* lead to worse patient outcomes.

### Pyoverdine production is induced by *P. aeruginosa* when cocultured with *Achromobacter* spp

During a previous screen of 1,597 bacterial isolates from pwCF for inhibitory activity (7), we observed that *Achromobacter* isolates increased production of a green pigment by *P. aeruginosa*. We repeated these coculture assays on synthetic cystic fibrosis sputum medium (SCFM) supplemented with 25 *µ*M ferric chloride (FeCl_3_) to mimic physiological concentrations of ferric iron (Fe^3+^) in the airways of pwCF (35, 36). We allowed the *Achromobacter* isolate to establish for 72 h after which we inoculate the *P. aeruginosa* perpendicularly to the established culture. After overnight coincubation, we confirmed that *Achromobacter dolens* strain RMS0405 induced production of a green pigment in *P. aeruginosa* strain PAO1 (**Fig. 2A**). We suspected that coculture with *A. dolens* induced pyoverdine, because we observed fluorescence when placing plates in a blue light (∼470 nm) transilluminator (**Fig. 2B**). We repeated cocultures with *A. dolens* using a Δ*pvdA* mutant that is unable to produce pyoverdine and confirmed loss of pigment production which was restored upon complementation with *pvdA* (Δ*pvdA* + *pvdA*) (**Fig. 2AB**). We observed no production of pyoverdine from *P. aeruginosa* in monoculture under these conditions (**Fig. 2C**). Together, these data indicate that *A. dolens* induces production of pyoverdine by *P. aeruginosa* during coculture.

**Figure 2.**
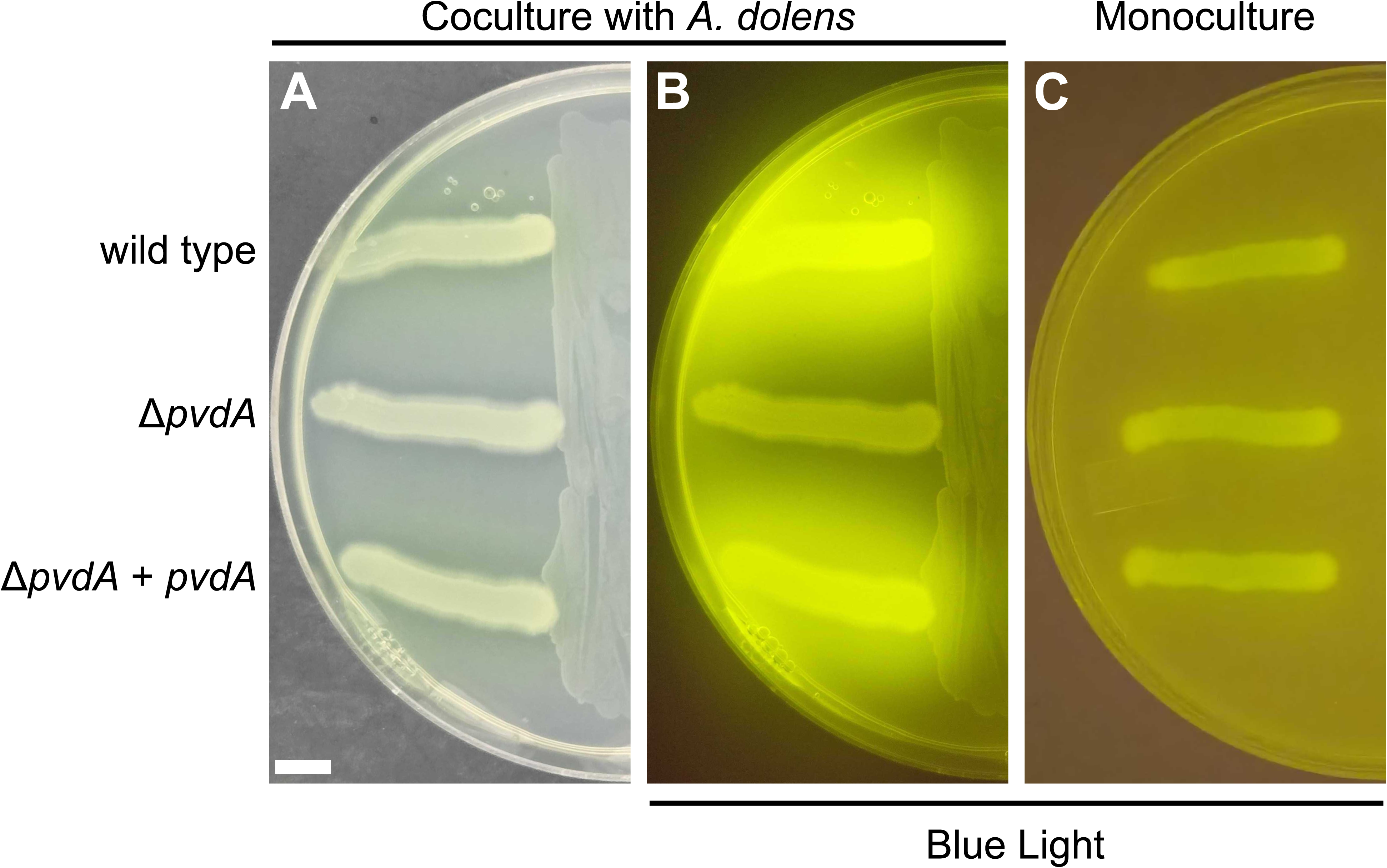
*A. dolens* induces pyoverdine production in *P. aeruginosa*. (**A**) Coculture of *A. dolens* RMS0405 (*Ad*, right) and *P. aeruginosa* strain PAO1 (left) on SCFM + 25 *µ*M FeCl_3_. Strains are labeled on the left: wild type (RMS0290), Δ*pvdA* (RMS0289), and Δ*pvdA* + *pvdA* (RMS0482). (**B**) Plate from panel **A** placed in a blue light transilluminator. Under blue light, pyoverdine exhibits fluorescence. (**C**) Cultures of *P. aeruginosa* growing on SCFM + 25 µM FeCl_3_ without *A. dolens* in a blue light transilluminator. All images are representative of triplicate samples. Scale bar is 5 mm.

### *A. dolens* induces pyoverdine production in clinical *Pseudomonas* isolates

Our assays were performed using *P. aeruginosa* strain PAO1, a laboratory strain that was originally isolated from an infected wound (37). To determine if pyoverdine induction extended beyond PAO1, we tested *A. dolens* strain RMS0405 against 48 clinical isolates of *Pseudomonas* from different pwCF that previously isolated (7) (**Table S2** in the supplemental material). Because most isolates were identified only by sequencing the 16S rRNA V3– V4 region, we refer to them as *Pseudomonas*. As a control, we also inoculated each of these *Pseudomonas* isolates on SCFM with no added iron (SCFM + 0 *µ*M FeCl_3_) without *A. dolens*, to assess whether they would produce pyoverdine under iron-limiting conditions.

Twenty (41.7%) isolates did not produce pyoverdine in monoculture or when cocultured. Accordingly, when we screened these isolates for *pvdA* using PCR, we found 5 (25%) did not yield a PCR product. We presume that the other 15 isolates possess mutations that abrogate pyoverdine biosynthesis. Two isolates (4.2%) produced pyoverdine in response to *A. dolens*, but not in response to iron limitation. However, eight (16.7%) isolates only produced pyoverdine in response to iron limitation. Finally, eighteen (37.5%) isolates produced pyoverdine in response to iron limitation and *A. dolens*.

Together, 18 of 26 (69.2%) *pvdA*-positive *Pseudomonas* isolates produced pyoverdine under iron-limiting conditions and during coculture, suggesting that the response to *A. dolens* is common, but not ubiquitous, in *Pseudomonas* (**Table 1**).

**Table 1.** *A. dolens* can induce pyoverdine production in clinical isolates of *Pseudomonas*.^1^ Pyoverdine production was scored by fluorescence after 24 h monoculture or coculture with *A. dolens* strain RMS0405. All isolates were screened in biological triplicate (independent experiments on different days). Nine isolates were inconsistent across the initial triplicates and the assays were repeated. On repeat, triplicates were consistent with each other and with the majority of replicates from the initial experiment. ^2^All isolates were screened by PCR using primers flanking *pvdA*. Isolates that were *pvdA*-negative were retested using genomic DNA extracted from a second culture to confirm absence of the PCR product.

| Pyoverdine Production<sup>1</sup> |  | *pvdA* PCR Product<sup>2</sup> |  |  |
| --- | --- | --- | --- | --- |
| Monoculture<br>(SCFM + 0 μM<br>FeCl<sub>3</sub>) | Coculture with *A.*<br>*dolens*<br>(SCFM + 25 μM FeCl<sub>3</sub>) | Yes | No | Total |
| No | No | 15 | 5 | 20 |
| No | Yes | 2 | 0 | 2 |
| Yes | No | 8 | 0 | 8 |
| Yes | Yes | 18 | 0 | 18 |
| Total |  | 43 | 5 | 48 |

### Multiple species of *Achromobacter* induce pyoverdine production in *P. aeruginosa*

We tested twenty additional strains of *Achromobacter* spp., encompassing seven of the eleven species previously reported in the airways of pwCF (12), isolated from multiple sources and tested their ability to induce pyoverdine production in *P. aeruginosa* strain PAO1, as above. In total, we found that 17/21 (81.0%) of *Achromobacter* spp. induced production of pyoverdine in *P. aeruginosa*. Each *Achromobacter* species that we tested was able to induce pyoverdine. However, 4/11 (36.4%) strains of *A. xylosoxidans* did not induce pyoverdine when cocultured with *P. aeruginosa* (**Fig. 3A**). We note here that strains DSM 6388, DSM 26073, and MN001 were deposited as *A. xylosoxidans*, but a core genome phylogeny and average nucleotide identity (ANI) to type species classified them as *Achromobacter insuavis* (99.2% ANI), *A. insuavis* (99.8% ANI), and *Achromobacter ruhlandii* (98.4% ANI), respectively. Together, these data suggest the ability to induce pyoverdine is widespread, but not universal, among *Achromobacter* spp.

**Figure 3.**
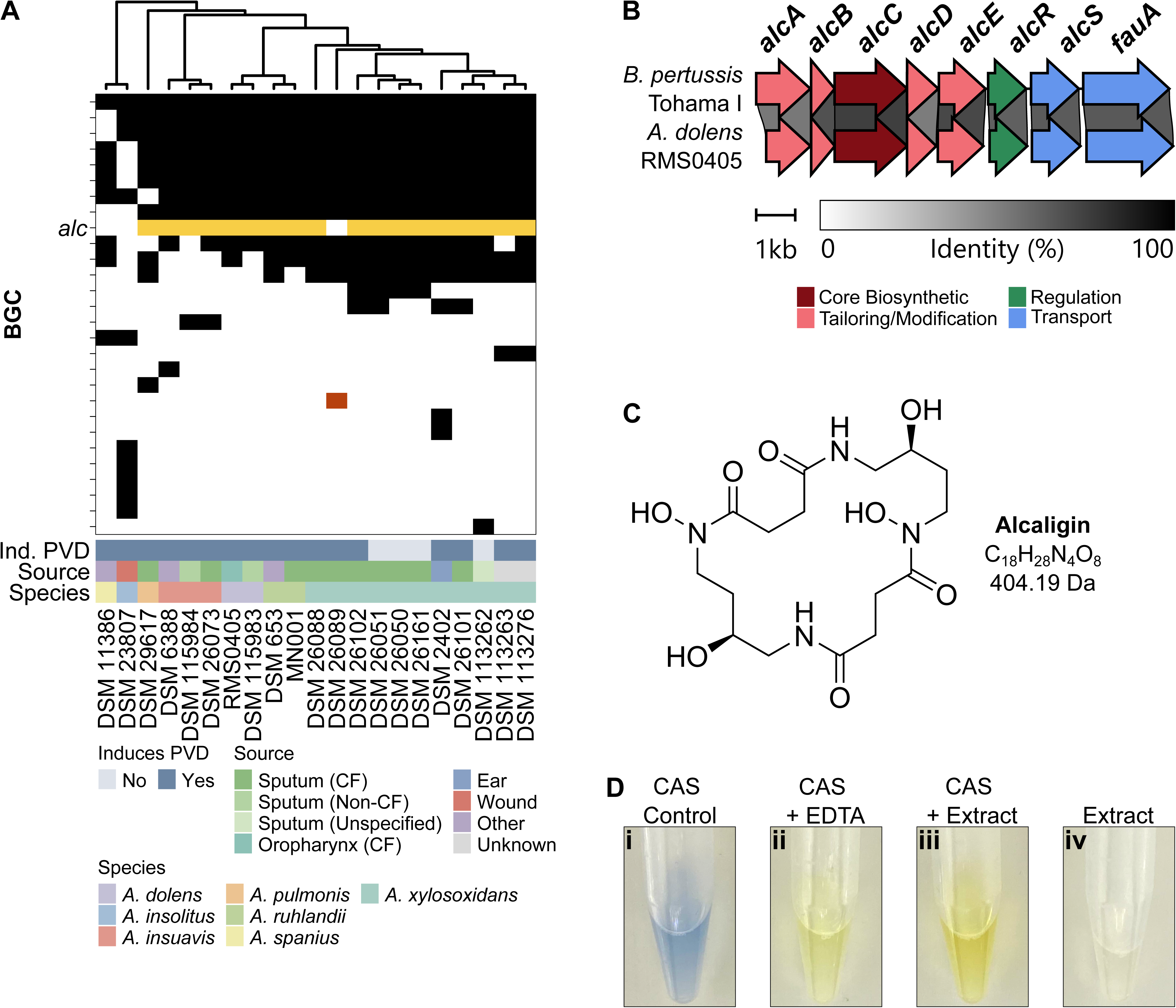
*Achromobacter* spp. encode BGCs for alcaligin and other secondary metabolites. (**A**) Heat map of BGCs identified within *Achromobacter* spp. genomes (horizontal), organized based on a core genome phylogeny (top). Presence and absence of a BGC (rows) indicated in black and white, respectively. The alcaligin (*alc*) BGC is highlighted in gold. A putative NRP-metallophore BGC is highlighted in rust. The key for *Achromobacter* spp. isolation source, species, and pyoverdine (PVD) induction is indicated below. (**B**) Comparison of the *alc* BGC from *A. dolens* strain RMS0405 and *B. pertussis* strain Tohama I. Darker shading corresponds to higher sequence identity, as indicated by the scale below. Gene color corresponds to protein function, as indicated by the key below. Scale bar is 1.0 kb. See **Fig. S2** in the supplemental material for comparisons to the corresponding BGCs from all *Achromobacter* spp. (**C**) Structure, formula, and predicted molecular mass of alcaligin. (**D**) CAS assay results from (i) a negative control, (ii) an ethylenediaminetetraacetic acid (EDTA) positive control, (iii) culture extract from *A. dolens* strain RMS0405. (iv) The culture extract without addition of CAS reagent.

### *Achromobacter* spp. encode a biosynthetic gene cluster for production of the siderophore alcaligin

Because pyoverdine is produced under iron limitation, we hypothesized that siderophores produced by *Achromobacter* may induce its production. Previous work demonstrated that siderophore production is widespread among *Achromobacter* isolates and is more common in clinical *Achromobacter* isolates than in environmental isolates (22). Therefore, to characterize siderophore production in *Achromobacter* spp., we performed whole genome sequencing for all *Achromobacter* strains that had not been sequenced, and used antiSMASH and BiG-SCAPE to identify biosynthetic gene clusters (BGCs) and group them into gene cluster families (GCFs).

Each *Achromobacter* spp. encoded a median of 12 BGCs or BGC fragments per genome (IQR: 11 to 13 BGCs or BGC fragments per genome) (**Fig. 3A**). Eighteen (86%) *Achromobacter* spp. strains, encompassing five of seven species that we tested, and all CF strains, encoded a BGC with high sequence similarity to the non-ribosomal independent siderophore (NIS) BGC for alcaligin (*alc*) from *Bordetella pertussis* strain Tohama I (**Fig. 3B** and **Fig. S2** in the supplemental material). Fourteen (82.3%) of *Achromobacter* strains that induced pyoverdine production encoded this BGC. Four strains of *A. xylosoxidans* encoded this BGC but did not induce pyoverdine. In addition, three *Achromobacter* spp. induced pyoverdine production, but did not encode the *alc* BGC (**Fig. 3A**).

Alcaligin (**Fig. 3C**) is a dihydroxamate iron-binding siderophore that is essential for lung colonization by *B. pertussis and Bordetella bronchiseptica* (38, 39). Alcaligin was originally reported from *A. xylosoxidans* strain KN 3-1 (formerly *Alcaligenes denitrificans* subsp. *xylosoxidans*), isolated from lagoon sediment (40, 41), but its production from CF isolates of *Achromobacter* has yet to be confirmed. We confirmed that all *Achromobacter* strains with the *alc* BGC possess all genes necessary for the proposed biosynthesis of alcaligin starting from L-ornithine (**Fig. S3A** in the supplemental material). These strains also encoded genes for the proteins involved in alcaligin regulation (*alcR*) (42–44), export (*alcS*) (45), uptake of ferri-alcaligin (*fauA*) (39, 46), and reductive release of iron (*alcD*) (43). There were weak hits to *alcS* and *fauA* in *alc* negative strains, which may represent other major facilitator superfamily or TonB-dependent transporters. As these strains lack *alcD*, it is unlikely that they would be able to utilize alcaligin (see **Fig. S2** and **Fig. S3B** in the supplemental material).

### *A. dolens* produces a siderophore

An agar extract from *A. dolens* strain RMS0405 cultured on SCFM possessed chrome azurol S activity (**Fig. 3D**), confirming siderophore production by this strain (47).

*A. dolens* produces alcaligin and *P. aeruginosa* produces pyoverdine and pyochelin during coculture.

We next examined siderophore production during coculture of *A. dolens* strain RMS0405 and *P. aeruginosa* strain PAO1 or the Δ*pvdA* mutant. Consistent with our previous results, we detected fluorescence from the wild-type strain (**Fig. 4A**), but not from the Δ*pvdA* mutant (**Fig. 4B**).

**Figure 4.**
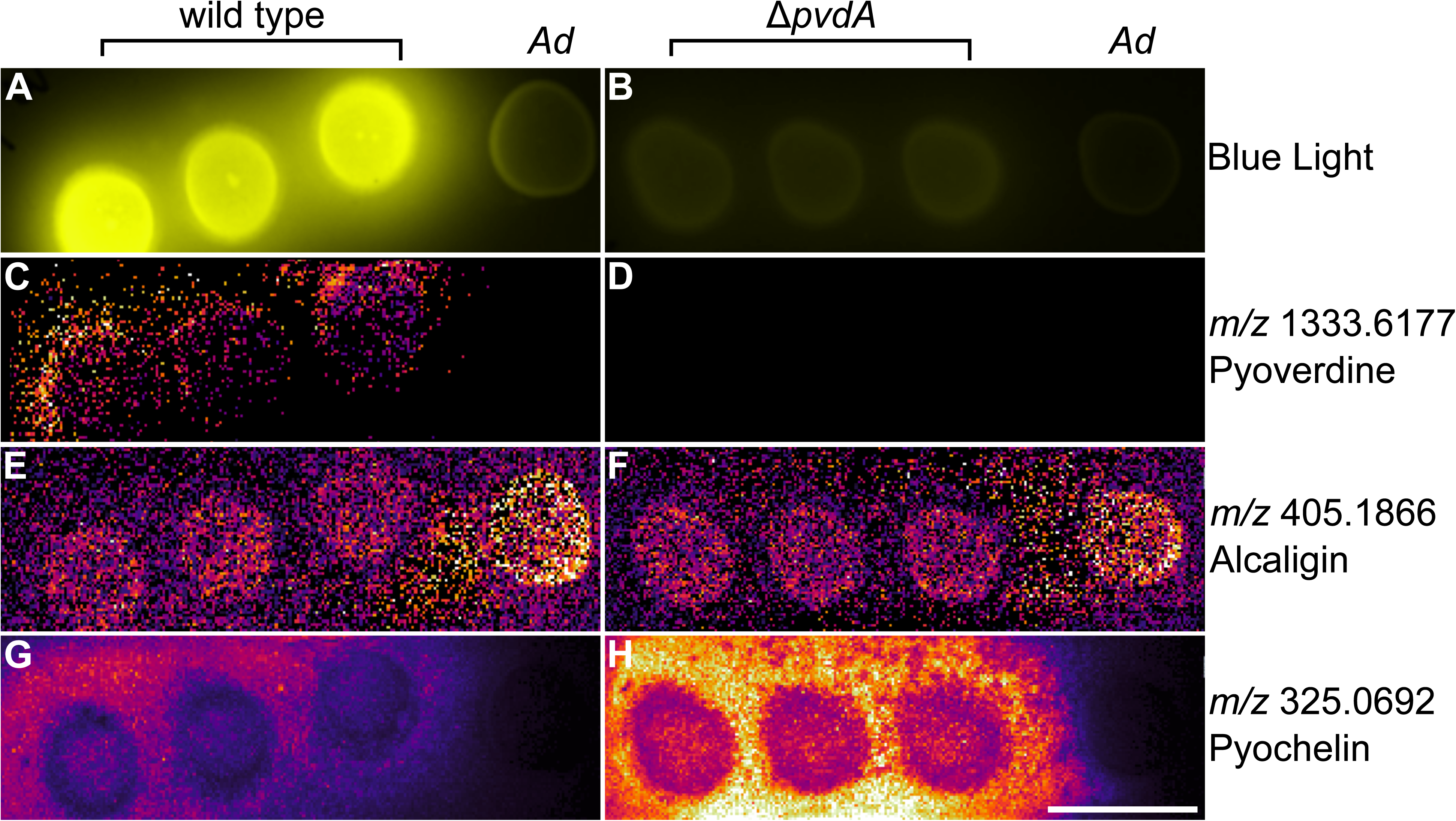
Production of Siderophores during coculture of *A. dolens* and *P. aeruginosa*. Mass spectrometry imaging of coculture of *A. dolens* strain RMS0405 (*Ad*, right) and *P. aeruginosa* strain PAO1 on SCFM + 25 *µ*M FeCl_3_. (**A-B**) False-colored blue light image of *P. aeruginosa* (**A**) wild type and (**B**) Δ*pvdA* and *A. dolens*. (**C-D**) Extracted ion images of molecular features ([M+H]^+^) for pyoverdine in *P. aeruginosa* (C) wild type and (**D**) Δ*pvdA*. (**E-F**) Extracted ion images of [M+H]^+^ for alcaligin in coculture between *P. aeruginosa* (**E**) wild type and (**F**) Δ*pvdA* and *A. dolens*. (**G-H**) Extracted ion images of [M+H]^+^ for pyochelin in coculture between *P. aeruginosa* (**G**) wild type and (**H**) Δ*pvdA* and *A. dolens*. Brighter pixels indicate higher signal of the corresponding [M+H]^+^. Note, the ion intensity for a given molecular feature is comparable between the cocultures with *P. aeruginosa* wild type (left images) and Δ*pvdA* (right images), but not between different ions due to differences in ionization efficiency. All *m/z* values are ± 4.6 parts per million (ppm). Scale bar is 5 mm. Experiment is representative of duplicate samples. See **Figs. S4** and **S5** in the supplemental material for extracted ion images of potential alcaligin analogues and known *P. aeruginosa* secondary metabolites, respectively.

We then used matrix-assisted laser desorption/ionization mass spectrometry (MALDI-MSI) to identify and map the distribution of metabolites produced by *A. dolens* and *P. aeruginosa* over the agar surface of the coculture. We identified an ion corresponding to pyoverdine E (*m/z* 1333.6177 ± 4.6 parts per million [ppm] [M+H]^+^), the major variant produced by *P. aeruginosa* strain PAO1 (48) from wild type (**Fig. 4C**), but not the Δ*pvdA* mutant (**Fig. 4D**). We then confirmed production of alcaligin (*m/z* 405.1866 [M+H]^+^) by *A. dolens* in both cocultures (**Fig. 4E-F**). We also identified two ions that likely correspond to alcaligin (*m/z* 405.1907 [M+H]^+^) and a +2 Da analogue (*m/z* 407.2865 [M+H]^+^) (**Fig. S4** in the supplemental material). We examined cocultures for production of pyochelin (*m/z* 325.0692 [M+H]^+^), the second major siderophore of *P. aeruginosa*. Strikingly, we observed that pyochelin was produced more abundantly by the Δ*pvdA* mutant than the wild type (**Fig. 4G-H**). We speculated that *P. aeruginosa* Δ*pvdA* increases production of pyochelin in coculture with *A. dolens* to compensate for loss of pyoverdine production. We generated a pyochelin biosynthesis mutant (Δ*pchA*) and a pyoverdine/pyochelin double biosynthesis mutant (Δ*pvdA* Δ*pchA*) and tested them in coculture with *A. dolens* on SCFM with no supplemented FeCl_3_. Under these conditions, we observed a minor growth defect of the Δ*pvdA* mutant that was more pronounced in the Δ*pvdA* Δ*pchA* mutant (**Fig. S5** in the supplemental material). We did not observe any other differences in production of known *P. aeruginosa* secondary metabolites between wild type or Δ*pvdA* (**Fig. S6** in the supplemental material).

We performed MALDI-QqTOF data-dependent tandem mass spectrometry (MALDI-TOF-DDA-MS^2^) to validate these siderophores. Based on a library match, we confirmed pyoverdine E (*m/z* 1333.6143) by the wild-type *P. aeruginosa* (**Fig. S7A** in the supplemental material). We assigned *m/z* 405.1980 as alcaligin because SIRIUS predicted the formula C_16_H_28_N_4_O_8_, consistent with alcaligin (**Fig. S7B** in the supplemental material), although the MALDI-MS/MS spectrum did not match predicted ESI fragmentation patterns in reference databases. This is likely due to difference in ionization sources and the collision energies that arise during ion formation. We have observed this phenomenon in other experiments, and this is an active area of research to determine how different ionization sources impact the resulting MS/MS spectra. Similarly, we assigned *m/z* 325.0675 as pyochelin based on an exact mass match to the expected formula and high-quality MS/MS data (**Fig. S7C** in the supplemental material). Together, these data indicate that siderophores are produced by both *A. dolens* and *P. aeruginosa* during coculture.

### Iron supplementation abolishes pyoverdine induction by *P. aeruginosa* during coculture with *A. dolens*

We hypothesized that competition for Fe^3+^ between *Achromobacter* spp. and *P. aeruginosa* leads to pyoverdine production. To test this hypothesis, we repeated the coculture assays between *A. dolens* strain RMS0405 and *P. aeruginosa* strain PAO1 on SCFM, but varied the concentration of supplemented Fe^3+^ (**Fig. 5**). With no added iron (outside of trace iron in our water and glassware), we observed production of pyoverdine in coculture (**Fig. 5A**) and monoculture (**Fig. 5D**). As before, on SCFM + 25 *µ*M FeCl_3_, pyoverdine was induced in coculture (**Fig. 5B**), but not monoculture (**Fig. 5E**). Finally, on SCFM + 100 *µ*M FeCl_3_, pyoverdine was not produced under either condition (**Fig. 5C, F**). For all experiments, there was no production of pyoverdine by the Δ*pvdA* mutant, and the complement behaved like wild type. These results support a model wherein pyoverdine is induced by iron depletion by *Achromobacter* spp. using alcaligin or other siderophores.

**Figure 5.**
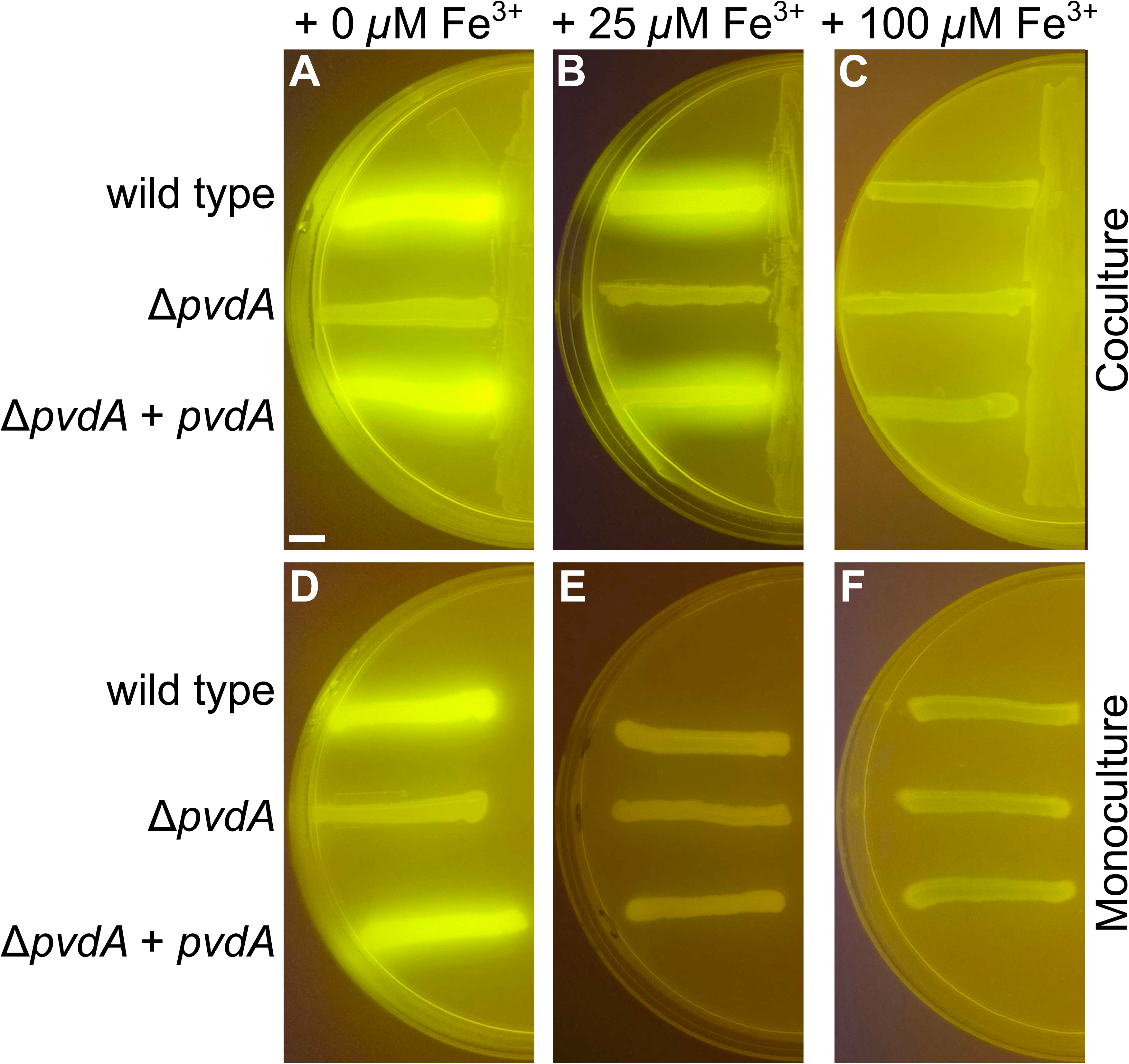
Iron modulates pyoverdine induction during *A. dolens*-*P. aeruginosa* cocultures. (**A-C**) Coculture of *A. dolens* strain RMS0405 (*Ad*, right) and *P. aeruginosa* strain PAO1 (left) on SCFM with (**A**) no supplemented FeCl_3_ (+ 0 *µ*M), (**B**) supplemented with 25 *µ*M FeCl_3_ (+ 25 *µ*M), and (**C**) supplemented with 100 *µ*M FeCl_3_ (+ 100 *µ*M). (**D-F**) Monoculture controls of *P. aeruginosa* on SCFM with (**D**) no supplemented FeCl_3_ (+ 0 *µ*M), (**E**) supplemented with 25 *µ*M FeCl_3_ (+ 25 *µ*M), and (**F**) supplemented with 100 *µ*M FeCl_3_ (+ 100 *µ*M). Strains are labeled on the left: wild type (RMS290), Δ*pvdA* (RMS0289), and Δ*pvdA* + *pvdA* (RMS0482). All images are representative of duplicate samples taken under a blue light transilluminator. Scale bar is 5 mm.

### Coculture with *A. dolens* activates expression of the low-iron stress response in *P. aeruginosa*

We compared the transcriptomes of *P. aeruginosa* cultured on SCFM + 25 *µ*M FeCl_3_ in monoculture and in coculture with *A. dolens*. We harvested RNA after 12 h, just before production of pyoverdine is visible under blue light. Coculture with *A. dolens* significantly altered the transcriptional profile of *P. aeruginosa*, resulting in 366 genes with increased expression and 253 genes with decreased expression (**Fig. 6A** and **Table S3** in the supplemental material). We observed that expression of most (14/16) pyoverdine biosynthesis genes (median: +1.5×, IQR: +1.4 to +1.6×, Benjamini-Hochberg [BH]-adjusted *P* = 3.e-15), and the *fpvA* (+1.3×, BH-adjusted *P =* 2.4e-2) and *fpvB* (+2.1×, BH-adjusted *P* = 1.0e-6) genes, which encode for ferripyoverdine importers, were increased during coculture. In addition, we found that expression other outer membrane TonB-dependent receptors (median: +1.3× IQR: +1.0 to +1.8×, BH-adjusted *P =* 2.5e-5), and the hemophore-encoding *hasAp* (49) (+1.6×, BH-adjusted *P =* 3.1e-2) were also increased during coculture.

**Figure 6.**
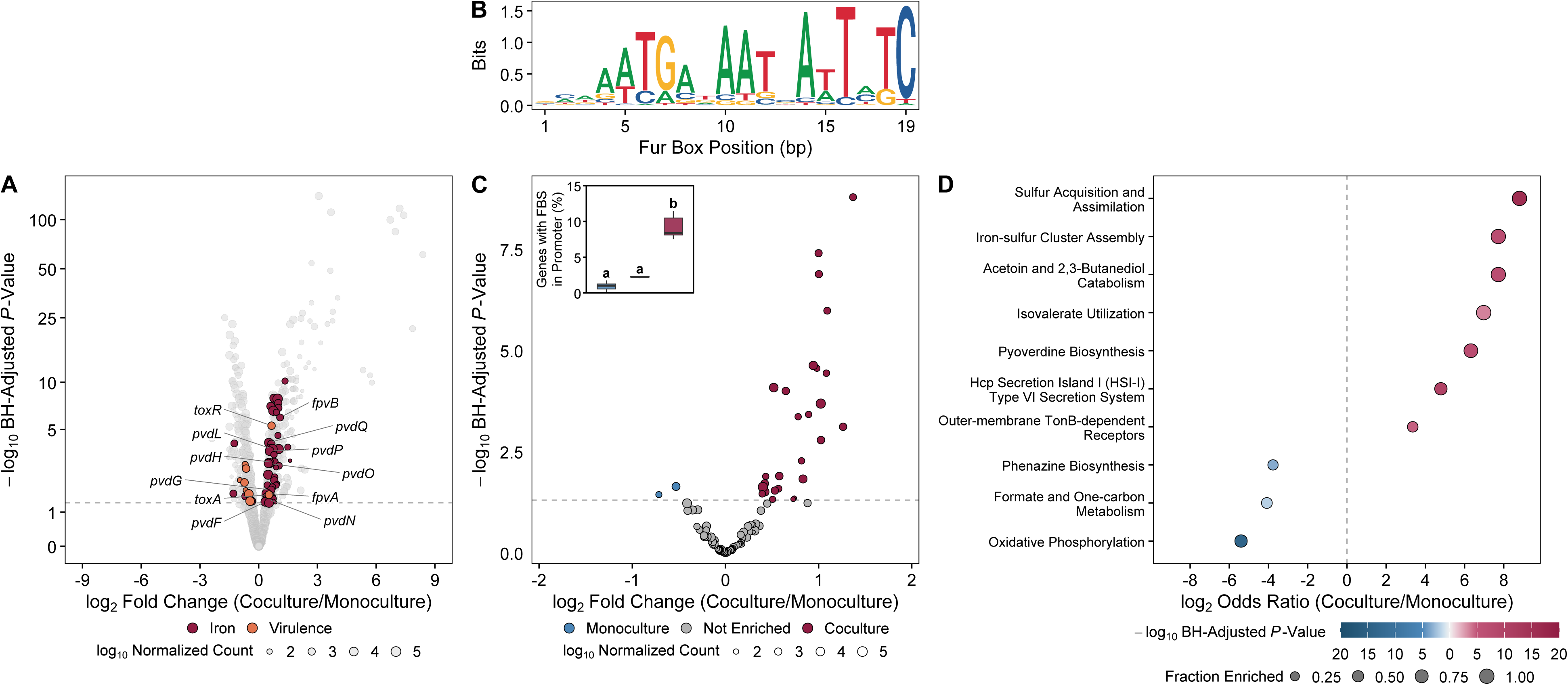
Coculture with *A. dolens* induces a low-iron stress response in *P. aeruginosa*. (**A**) Volcano plot showing the fold change of genes from *P. aeruginosa* strain PAO1 that were detected in SCFM supplemented with 25 *µ*M FeCl_3_ either in monoculture or coculture with *A. dolens* strain RMS0405. Each point represents a gene, and its size corresponds to the normalized read count, as indicated below. The color of the points indicates protein functional category, as indicated below. The horizontal dashed line indicates the BH-adjusted *P*-value threshold of 0.05 for significance. The genes involved in pyoverdine biosynthesis and exotoxin production and regulated are highlighted. (**B**) Sequence logo for 105 predicted FBS sequences in the *P. aeruginosa* strain PAO1 genome. (**C**) Subset of the volcano plot in **A** only showing genes with a predicted FBS. Inset: box plots of the percentage of *P. aeruginosa* strain PAO1 genes with putative FBS with increased expression in monoculture (blue), with increased expression in coculture with *A. dolens* strain RMS0405 (red), or were not differentially regulated (grey). Each box summarizes eight results obtained by withholding one sample and repeating the differential expression analysis. The upper and lower bounds of the box plots indicate the 75th and 25th percentiles, respectively. The horizontal black bar indicates the median. Conditions that share a letter are not significantly different (BH-adjusted *P* > 0.05). (**D**) Plot showing *P. aeruginosa* strain PAO1 pathways increased or decreased during coculture with *A. dolens* strain RMS0405. The color and size of each point represents the −log_10_ BH-adjusted *P* value and the fraction of genes differentially regulated, respectively, as indicated below.

We identified a total of 95 promoters in the *P. aeruginosa* strain PAO1 genome that contained putative ferric uptake regulator (Fur) binding site (FBS) sequences (**Fig. 6B** and **Table S4** in the supplemental material). We found that 28 (29.5%) of these genes were upregulated during coculture, compared to 338/5,333 (6.3%) of genes lacking an upstream putative FBS (Fisher’s exact OR = 6.2, *P* = 8.4e-12) (**Fig. 6C**). 10.4% of genes with increased coculture expression had putative FBS, which was a 9.0-fold increase over monoculture (Fisher’s exact odds ratio [OR] = 9.8, *P* = 9.9e-5) (**Fig. 6C inset**).

There were 10 gene sets with increased (n=7) or decreased (n=3) expression in *P. aeruginosa* during coculture (**Fig. 6D**). These decreased pathways encode proteins that are large iron sinks. For instance, the genes for NADH dehydrogenase I (complex I) and the cytochrome bc_1_ complex in the “oxidative phosphorylation” pathway use Fe-S clusters and a Rieske [2Fe-2S] center together with heme, respectively, are decreased. In addition, the genes for *fumC1* (+2.0×, BH-adjusted *P =* 5.7e-3), which encodes an alternative non-Fe-S-containing fumarase and *mqoA* (+2.4×, BH-adjusted *P =* 2.9e-8), which encodes a malate:quinone oxidoreductase that bypasses complex I, are upregulated in during coculture. Within the “formate and one-carbon metabolism pathway”, expression of the catalase *katA* (−2.8×, BH-adjusted *P =* 2.5e-8) and formate dehydrogenase *fdnGHI* (−1.5 to −1.7×, BH-adjusted *P* ≤ 1.0e-2) genes, which contain heme and Fe-S clusters, respectively, are decreased. Furthermore, expression of *acnA* (−1.4×, BH-adjusted *P =* 2.1e-3), which encodes aconitase, and *moaA1* (−2.8×, BH-adjusted *P =* 5.8e-4) and *moaA2* (−1.4×, BH-adjusted *P =* 1.3e-2) genes for molybdopterin biosynthesis, which each harbor two [4Fe-4S] clusters were decreased. Moreover, the *isc* operon for Fe-S cluster assembly was increased (+1.3 to +1.8×, BH-adjusted *P* = 1.3e-12), potentially due lack of a [2Fe-2S] cluster in the regulator IscR. Together, these data suggest that coculture with *A. dolens* stimulates a low-iron stress response in *P. aeruginosa*.

### Coculture with *A. dolens* may also shift *P. aeruginosa* to alternative carbon and sulfur sources and alter expression of type VI secretion and phenazine biosynthesis genes

Coculture with *A. dolens* strain RMS0405 significantly altered expression of metabolic pathways in *P. aeruginosa* strain PAO1 (**Fig. 6D**). In coculture, *P. aeruginosa* upregulated expression of the *bkdA1*, *bkdA2*, and *bkdB* (+1.4× to +1.7×, BH-adjusted *P* < 1.0e-3) and *liuABCD* (median: +1.8×, IQR: +1.7 to +1.9×, BH-adjusted *P =* 2.9e-4) genes, which are involved in leucine catabolism. This suggests that *A. dolens* depletes the available glucose and lactate in SCFM before *P. aeruginosa* is cultured. The most highly induced genes during coculture were for acetoin and 2,3-butanediol metabolism (median: +150×, IQR: +120 to +200×, BH-adjusted *P =* 1.2e-7). Accordingly, *A. dolens* encodes a diacetyl reductase that is predicted to produce (*S*)-acetoin as a byproduct of pyruvate metabolism. Furthermore, *A. dolens* may deplete the available sulfur, as all 15 genes for sulfate transport and assimilation and for taurine and alkanesulfonate scavenging were upregulated (median: +2.6×, IQR: +1.9 to +3.4×, BH-adjusted *P =* 2.1e-16) during coculture.

Coculture with *A. dolens* increased expression of the Hcp secretion island I (H1) type VI secretion system (T6SS) genes (median: +1.6×, IQR: +1.3 to +1.7×, BH-adjusted *P =* 1.2e-10) in *P. aeruginosa* (**Fig. 6D**). The T6SS is involved in interspecies bacterial competition and delivers toxic effectors into the periplasm of target cells (50). Expression of the effector genes *tse1* (+1.8×, BH-adjusted *P =* 8.7e-4) and *tse3* (+1.5×, BH-adjusted *P =* 6.4e-4) increased in coculture. Coculture also affected expression of phenazine biosynthesis genes. Expression of the *phzA2* (−1.6×, BH-adjusted *P =* 1.6e-4), *phzB2* (−2.3×, BH-adjusted *P =* 7.5e-10), and *phzH* (−2.0×, BH-adjusted *P =* 4.5e-6) genes decreased, while *phzA1* (+1.4×, BH-adjusted *P =* 1.2e-2) of the duplicate operon increased. However, we did not observe differences in the production of pyocyanin (*m/z* 211.086 [M+H]), phenazine-1-carboxylic acid (*m/z* 225.0671 [M+H]), phenazine dicarboxylic acid (*m/z* 269.1755), or phenazine-1-carboxamide (*m/z* 224.0824) between *P. aeruginosa* wild type and Δ*pvdA* during coculture with *A. dolens* (**Fig. S6G-N**). Together, these data suggest that coculture with *A. dolens* may alter the carbon and sulfur sources used by *P. aeruginosa,* and affect production of the T6SS.

### *A. dolens* stimulates production of extracellular virulence factors by *P. aeruginosa*

Pyoverdine positively regulates production of the virulence factors exotoxin A (ToxA) and the PrpL serine protease in *P. aeruginosa* acting through the FpvA-FpvR-PvdS signaling cascade (51, 52). We hypothesized that coculture with *A. dolens* may also stimulate production of these virulence factors. Expression of the *toxA* (+1.4×, BH-adjusted *P =* 2.6e-2) and *toxR* (+1.6×, BH-adjusted *P =* 5.1e-6) regulator genes increased during coculture (**Fig. 6A**). However, the expression of *prpL* was unchanged (+1.10×, BH-adjusted *P =* 0.64).

We wanted to verify if these transcriptional changes corresponded to changes in the production of ToxA protein. We excised the agar near *P. aeruginosa* growing next to *A. dolens*, and extracted proteins. We then used untargeted liquid chromatography-tandem mass spectrometry (LC-MS^2^) to identify these proteins. We detected 65 extracellular proteins. Note, though we took care not to disturb the *P. aeruginosa* streak when we excised the agar, we also detected cytoplasmic (n=1,451), cytoplasmic membrane (n=443), periplasmic (n=143), outer membrane (n=89), flagella (n=3), and proteins with unknown localization (n=449), which are likely derived from cell lysis (**Table S6** in the supplemental material). Across 2,633 proteins we detected in agar whose corresponding genes we also detected with transcriptomics, we noted a weak correlation between the protein abundance and transcriptional fold changes (Spearman ρ = 0.18, *P* = 1.8e-21).

In total, we identified 19 extracellular proteins that were present at higher levels during coculture (**Fig. 7**). Specifically, production and/or secretion of extracellular virulence factors was increased in coculture (Fisher’s exact OR: 11.0, *P* = 3.4e-5). These virulence factors included AprA (+4.9×, BH-adjusted *P =* 1.3e-3), LasB (+2.7×, BH-adjusted *P =* 1.2e-3), PaaP (+2.7×, BH-adjusted *P =*5.7e-3), PlcB (+2.2×, BH-adjusted *P =* 1.3e-2), and ToxA (+2.2×, BH-adjusted *P =* 3.4e-3). In addition, the hemophore HasAp (+4.4×, BH-adjusted *P =* 6.7e-3), which is known to be induced under iron-limiting conditions (53), was present at higher levels during coculture. Three extracellular proteins were decreased in coculture, including the S4 pyocin (−3.0×, BH-adjusted *P =* 1.2e-2). Taken together, our data indicate that *A. dolens* modulates multiple iron-responsive pathways in *P. aeruginosa* and stimulates production of virulence factors.

**Figure 7.**
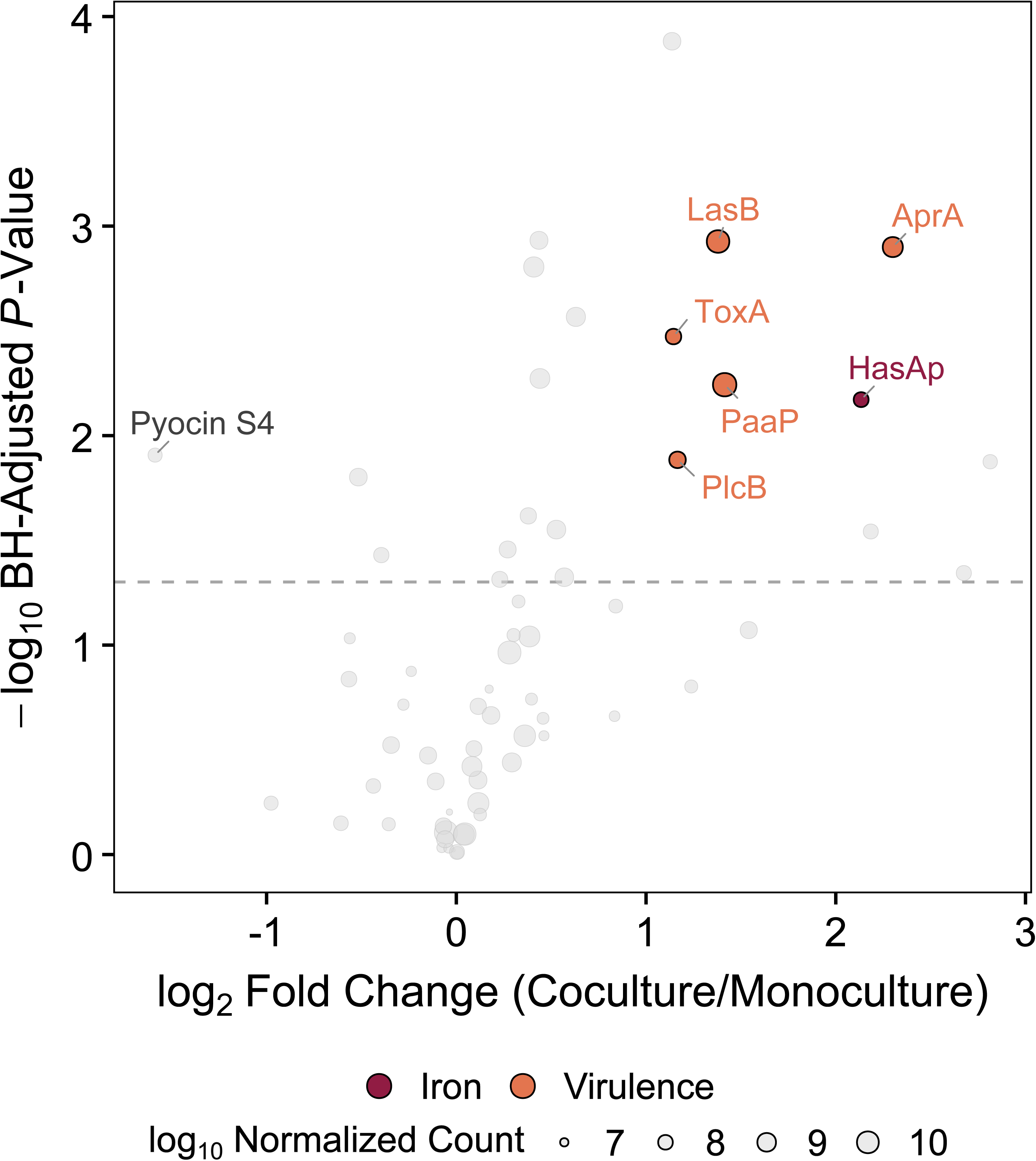
Coculture with *A. dolens* stimulates virulence factor production by *P. aeruginosa*. Volcano plot showing the fold change of proteins from *P. aeruginosa* strain PAO1 that were detected in SCFM supplemented with 25 *µ*M FeCl_3_ either in monoculture or coculture with *A. dolens* strain RMS0405. Each point represents a protein, and its size corresponds to the normalized protein intensity, as indicated below. The color of the points indicates protein functional category, as indicated below. The horizontal dashed line indicates the BH-adjusted *P*-value threshold of 0.05 for significance. Specific virulence factors and the hemophore HasAp are highlighted.

## Discussion

In this study we investigated interactions between the CF pathogens *Achromobacter* spp. and *P. aeruginosa*. Using existing sequencing data, we confirmed that *Achromobacter*-*P. aeruginosa* coinfections are associated with more severe loss of lung function than infection with either of these pathogens alone. Critically, this effect was specific to *Achromobacter*, as coinfections with *P. aeruginosa* and other CF bacterial pathogens did not exacerbate lung function decline (**Fig. 1**). We then determined that *Achromobacter* spp. that are known to infect the airways of pwCF induced production of the siderophore and virulence factor pyoverdine when cocultured with *P. aeruginosa* (**Fig. 2** and **Fig. 3**). We found that this phenomenon occurred broadly, as *Achromobacter* induced pyoverdine in multiple clinical *Pseudomonas* isolates from pwCF (**Table 1**). We sequenced the genomes of the *Achromobacter* spp., and we tested and determined that most contained a BGC for the siderophore alcaligin (**Fig. 3**). We confirmed that during coculture *A. dolens* produces alcaligin and *P. aeruginosa* produces the siderophores pyoverdine and pyochelin (**Fig. 4**). We then determined that pyoverdine induction in coculture was dependent on iron (**Fig. 5**). Finally, we found that coculture with *A. dolens* stimulates expression and production of multiple virulence factors by *P. aeruginosa* (**Fig. 6** and **Fig. 7**).

Within the environment of the CF airway, iron is available via multiple sources. For instance, hemorrhage introduces heme into the airway, and proteolysis of transferrin and lactoferrin may release ferric iron (54). Furthermore, cultured airway epithelial cells that express ΔF508-CFTR release more iron into the apical medium than cells that express a fully functional CFTR (55). As one means to acquire this iron, microbes produce siderophores. Indeed, 17 siderophores from the CF airway microbiota have been reported, making them one of the most commonly described classes of secondary metabolites from CF microbes to date (56). Siderophore production is required for many pathogens to colonize and infect the airway. For example, *B. pertussis fauA* mutants exhibit defects in murine lung colonization due to their inability to import ferric alcaligin, and *B. bronchiseptica* alcaligin biosynthesis mutants display broad colonization defects across the swine respiratory tract and cause less severe disease (38, 39). In the case of *P. aeruginosa*, production of pyoverdine has been confirmed in the lungs of pwCF (57, 58) and supports airway colonization (59–61). However, reliance on siderophores by *P. aeruginosa* changes over the course of infection. Chronic infection isolates often harbor mutations in pyoverdine biosynthesis as the source of iron in the CF airway changes (62). How changes in iron availability and siderophore production influence competition between CF pathogens over the course of chronic infection remains poorly understood, but our data suggest that these interactions may have important consequences for disease.

Ferripyoverdine signaling through the FpvA-FpvR-PvdS signaling cascade promotes production of pyoverdine and the virulence factors exotoxin A and PrpL (52). Unexpectedly, we did not observe increased production of PrpL in *P. aeruginosa* during coculture with *A. dolens*. However, coculture with *A. dolens* increased production of four additional secreted virulence factors (AprA, LasB, PaaP, and PlcP) (**Fig. 7**). There are no FBS sequences upstream of the corresponding genes for these virulence factors. We currently do not know how *A. dolens* induces their production. However, with the exception of *toxA*, none of these genes were upregulated, suggesting post-transcriptional regulation.

Pyoverdine itself is cytotoxic toward macrophages, kills *Caenorhabditis elegans* directly, and increases *P. aeruginosa* virulence in murine burn wound, endocarditis, and pneumonia models (63–67). Consequently, under iron-limitation, *P. aeruginosa* becomes more virulent (68). Therefore, we hypothesize that the lung function decline exacerbated during *Achromobacter*-*P. aeruginosa* coinfections occur, in part, as a byproduct of exploitation competition for iron between these pathogens. Notably, while alcaligin is a high-affinity iron chelator (−log_10_[Fe^3+^]_Free_ when [Fe^3+^] = 1 *µ*M, [Siderophore] = 10 *µ*M, pH 7.4 [pFe]: 23), pyoverdine is four orders of magnitude stronger (pFe: 27) (41, 69). This does not support a simple model in which alcaligin sequesters all available iron during competition and completely starves *P. aeruginosa* of iron. Total iron starvation is likely not required to induce siderophore production in *P. aeruginosa* as a response. In support of this idea, pyochelin (pFe: 16) (70) is a substantially weaker siderophore than both alcaligin and pyoverdine. We detected increased pyochelin production by Δ*pvdA* (compare **Fig. 4G** and **Fig. 4H**). If *A. dolens* depleted all available iron, then we would expect complete growth inhibition of the Δ*pvdA* Δ*pchA* double mutant (**Fig. S5**). Instead, our data suggest that iron limitation imposed in our coculture conditions is sufficient to alter *P. aeruginosa* siderophore production without fully preventing growth. Within the spatially structured environment of the CF airway, alcaligin could reduce local iron concentration sufficiently to induce pyoverdine (71–73). It is tempting to speculate that increased severity of *Achromobacter*-*Pseudomonas* coinfections is due solely to production of alcaligin as a relatively strong siderophore. However, the relative strengths of siderophores produced by CF isolates *Burkholderia* spp., *S. aureus*, and *Stenotrophomonas maltophilia* are not reported (74–81), making these comparisons impossible at this time.

Previous clinical studies (12) and the data we present suggest that *Achromobacter* increases *P. aeruginosa* virulence. However, the opposite effect on virulence in zebrafish was recently reported (29). Notably, these effects are strain-dependent, as Besse and Menetrey et al. found that strain Ax 200 reduced virulence, while the clonal strain Ax 198, isolated from the same patient, did not. Intriguingly, these authors also report that production of the pyochelin receptor FptA by *P. aeruginosa* strain II.17 decreased during coculture with *A. xylosoxidans* strain Ax 200 and Fur decreased during coculture with either strain Ax 198 or Ax 200. Similarly, pyoverdine production by II.17 was reduced during coculture with either *A. xylosoxidans*. Together, these data led Besse and Menetry et al. to propose that iron uptake by *P. aeruginosa* might be inhibited by *A. xylosoxidans* (29). These findings contrast with what we observed. It is not known if strains Ax 198 or Ax 200 produce alcaligin, but they do encode the *alc* BGC. We do note that four *A. xylosoxidans* strains we tested encode the *alc* BGC, but did not induce pyoverdine (**Fig. 3A**). It is possible that *A. xylosoxidans* strains Ax 198 and Ax 200, do not produce alcaligin. Regardless, the results of our experiments (**Figs. 6** and **7**), provide support for the idea that iron homeostasis is affected in *P. aeruginosa* during coculture with *A. dolens*.

Currently, our data suggest that alcaligin-imposed iron limitation induces pyoverdine production, but we have not directly demonstrated this mechanism. Thus far, our attempts to generate an alcaligin biosynthesis mutant in *A*. *dolens* strain RMS0405 have not been successful. Further, four strains of *A. xylosoxidans* encode *alc*, but do not induce pyoverdine in cocultured (**Fig. 3A**). We hypothesize that this discrepancy may be due to lack of alcaligin production. However, we have not identified any specific mutations in these strains that we would expect to impact alcaligin. In contrast, we observed that *Achromobacter* strain DSM 23807, *Achromobacter spanius* strain DSM 11386, and *A. xylosoxidans* strain DSM 26089 induce pyoverdine, but lack the *alc* BGC (**Fig. 3A**). In the case of *A. xylosoxidans* strain DSM 26089, we identified a BGC for a predicted non-ribosomal peptide metallophore, which is currently under investigation. However, we did not identify siderophore BGCs in *A. insolitus* or *A. spanius*. Perhaps these strains encode siderophores that we have not detected or there are alternative pathways to pyoverdine induction that remain to be discovered.

We last note an intriguing pattern we observed. The H1 T6SS genes had increased expression in *P. aeruginosa* during coculture, and this system functions broadly in interspecies bacterial competition (82). Conversely, the protein whose abundance decreased the most in coculture was pyocin S4. This protein resembles a colicin and that uses FpvA as its receptor for intraspecies competition (83). We speculate that coculture with *A. dolens* could cause *P. aeruginosa* to repress pyocin S4 as expression of *fpvA* increases, shifting this pathogen from intraspecies to interspecies competition, as evidenced by increased expression of the H1 T6SS genes. Subsequent work will be required to investigate if coculture with *A. dolens* alters competitive phenotypes broadly in *P. aeruginosa*.

In conclusion, we report that *Achromobacter* spp. induce production of pyoverdine by *P. aeruginosa* during coculture. Pyoverdine is itself a virulence factor that also regulates production of other virulence factors. Our observation may serve to explain the basis underlying the increased severity of disease in pwCF that are coinfected with both of these pathogens. It is of note that this phenomenon is not limited to *Achromobacter*, as Herzog et al. recently demonstrated that another group of emerging CF pathogens, the *Pandoraea* spp., produce siderophores called pandorabactins that induce pyoverdine production in *P. aeruginosa* (84). Thus, the potential for increased virulence of *P. aeruginosa* in coinfections may be more common than previously appreciated.

## Materials and Methods

### Achromobacter and Pseudomonas coinfection analyses

We previously described a reanalysis of 16S rRNA gene amplicon sequencing studies from CF airway samples (7). In the present study, we used these same data to assess the effects of coinfection with *Achromobacter* and *Pseudomonas* on lung function. Using a 1% relative abundance cutoff, we assigned samples to the following groups: no pathogen present, *Pseudomonas* only, *Achromobacter* only, or both pathogens. To confirm that the coinfection association was robust to the threshold used to define presence, we recomputed the difference in mean FEV_1_ pp values between samples with both pathogens and samples without *Pseudomonas* nor *Achromobacter* across thresholds of 0.1, 0.5, 1, 2, and 5% relative abundance, and estimated each mean difference and its 95% CI with Welch’s *t*-test. We excluded samples that lacked both *Pseudomonas* and *Achromobacter* but contained at least one other CF pathogen (*Burkholderia*, *Haemophilus*, *Staphylococcus*, or *Stenotrophomonas*), so that the no pathogen group was free of all six CF pathogens. We compared FEV_1_ pp values among the groups using pairwise Wilcoxon rank-sum tests and adjusted *P*-values for multiple comparisons with the Benjamini-Hochberg (BH) false discovery rate. We defined severe disease as samples where the FEV1 pp value was ≤30%. To compare the burden of severe disease for each group, we estimated the proportion of severe samples and the 95% confidence interval (CI) with the exact binomial method. We compared each group to the both pathogens group using Fisher’s exact test and adjusted the resulting *P*-values with the BH method.

To test whether lung function decline scaled with the degree of coinfection with *Achromobacter* and *Pseudomonas*, we related the combined relative abundance of *Achromobacter* and *Pseudomonas* to the FEV_1_ pp value using Spearman’s rank correlation. To test whether the association with worse lung function was specific to *Achromobacter*, we restricted the data to *Pseudomonas*-positive samples and fit a single multivariable ordinary least-squares model, FEV_1_ pp ∼ *Achromobacter* + *Burkholderia* + *Haemophilus* + *Staphylococcus* + *Stenotrophomonas* + Age + BioProject Accession, in which each pathogen term was a binary indicator of presence (≥1% relative abundance). We interpreted each pathogen coefficient as the adjusted change in FEV_1_ pp value associated with copresence of that pathogen, relative to *Pseudomonas*-positive samples lacking that pathogen and with the remaining pathogens held constant, and we report 95% CIs and per-coefficient *t*-test BH-adjusted *P*-values. We restricted this model to samples with complete covariate data.

### Bacterial strains and media

The *Pseudomonas* isolates and *Achromobacter* strains used in this study are provided in **Tables S2 and S7**, respectively, in the supplemental material. For general propagation and manipulation, we used lysogeny broth (LB; 1.0% [wt/vol] tryptone, 0.5% [wt/vol] yeast extract, 0.5% [wt/vol] sodium chloride, 1 mM sodium hydroxide). All plates were solidified using 1.5% (wt/vol) agar.

For coculture assays, we used SCFM (1.3 mM sodium dihydrogen phosphate, 1.25 mM disodium hydrogen phosphate, 0.35 mM potassium nitrate, 0.27 mM potassium sulfate, 2.3 mM ammonium chloride, 15 mM potassium chloride, 52 mM sodium chloride, 10 mM 3-(N-morpholino)propanesulfonic acid, 1.4 mM serine, 1.5 mM glutamate, 1.7 mM proline, 1.2 mM glycine, 1.8 mM alanine, 1.1 mM valine, 0.6 mM methionine, 1.1 mM isoleucine, 1.6 mM leucine, 0.7 mM ornithine, 2.1 mM lysine, 0.3 mM arginine, 0.01 mM tryptophan, 0.8 mM aspartate, 0.8 mM tyrosine, 1 mM threonine, 0.2 mM cysteine, 0.5 mM phenylalanine, 0.5 mM histidine, 3 mM glucose, 9 mM lactate, 0.6 mM magnesium chloride, 1.7 mM calcium chloride, 3.6 *µ*M iron (II) sulfate, pH 6.8) (85). To make SCFM agar plates, we mixed 3% [wt/vol] agar with double-strength SCFM. We selected exconjugants during complementation of *P. aeruginosa* mutants using Vogel-Bonner minimal medium (VBMM; 0.02% [wt/vol] magnesium sulfate heptahydrate, 1% [wt/vol] dipotassium hydrogen phosphate, 0.2% [wt/vol] citric acid, 0.35% [wt/vol] ammonium sodium phosphate dibasic) agar plates containing tetracycline.

### Identification of *Achromobacter* isolates

We isolated *A. dolens* strain RMS0405 as part of a previous study (7). Briefly, we used MacConkey agar (1.7% [wt/vol] gelatin peptone, 1.0% [wt/vol], sodium chloride, 0.15% [wt/vol] bile salts, 0.15% [wt/vol] casein peptone, 0.15% [wt/vol] meat peptone, 104 *µ*M neutral red, 2.5 *µ*M crystal violet, 1.35% [wt/vol] agar, pH 7.1) supplemented with 0.5% [wt/vol] xylose with the following additions after autoclaving: 20 *µ*g/mL vancomycin, 20 *µ*g/mL aztreonam, and 5 *µ*g/mL amphotericin B (MCXVAA). We cultured bacterial strains in 3 mL of BHI (1.45% [wt/vol] casein peptone, 0.7% [wt/vol] meat peptone, 0.6% [wt/vol] brain heart infusion solids, 0.5% [wt/vol] sodium chloride, 0.25% [wt/vol] disodium phosphate, 0.2% [wt/vol] dextrose) in a shaker at 37°C. We extracted DNA from the overnight liquid culture using the phenol-chloroform extraction method. We amplified the full-length 16S rRNA gene using the universal primers 27F (5’-AGAGTTTGATCMTGGCTCAG-3’) and 1492R (5’-CGGTTACCTTGTTACGACTT-3’) and performed Sanger sequencing at the Oklahoma State University Genomics and Proteomics Center.

### Cross-streak coculture assays

We inoculated 3 mL LB with single colonies of *Achromobacter* spp. isolates and incubated the cultures at 37°C overnight with shaking. We then adjusted the optical density at 600 nm (OD_600_) of the overnight culture to 1.0 and spread 50 *µ*L in a ∼8 × ∼1.5 cm-wide line in the center of a 25 mL SCFM agar plate (diameter: 8.6 cm). We incubated the plates for 72 h at 37°C. We prepared overnight (16 to 18 h) cultures of *P. aeruginosa* strains by inoculating single colonies into 3 mL LB. We diluted the overnight cultures to OD_600_ = 0.01. We then used sterilized wooden applicator sticks to streak the cultures perpendicularly to the *Achromobacter* spp. streak. We returned the plates to 37°C and assessed pyoverdine production after overnight incubation under blue light (λ = 470 nm) using a transilluminator.

### Genomic DNA extraction, sequencing, and assembly

We extracted genomic DNA from *Achromobacter* spp. using previously reported method (86). Briefly, we washed cell pellets from 3 ml of an overnight culture in BHI broth in 1 ml of Buffer 1 (150 mM NaCl, 10 mM EDTA, 20 mM Tris-HCl, pH 8). We then resuspended the cells in 1 mL of fresh Buffer 1 with 1 mg RNase A and SDS to a final concentration of 1.4% (wt/vol). We incubated the samples at 37°C for 90 min. We then added 0.4 mg of proteinase K and incubated samples at 65°C for 20 min. We incubated samples at 37°C for 16 h to ensure complete lysis. Subsequently, we extracted and purified genomic DNA used standard phenol-chloroform extraction and followed by isopropanol precipitation.

We submitted genomic DNA from *A. dolens* strain RMS0405 and *A. xylosoxidans* DSM 26089 to Plasmidsaurus for whole genome sequencing using Oxford Nanopore Technology. Libraries were prepared using v14 library prep chemistry and sequenced using R10.4.1 flow cells. Bases were called using Dorado v4.3 with default Q10 quality filtering (https://github.com/nanoporetech/dorado). Then Filtlong v0.2.1 (https://github.com/rrwick/Filtlong) was used to remove reads <300 bp and retain the 95% highest quality reads. Subsequently, Autocycler (87) was used to generate assemblies using Flye v2.9.6+ (88) and Hifiasm (89), remove low depth and small contigs, then compress, cluster, trim and, resolve the assemblies. For all other *Achromobacter* spp., we submitted genomes to SeqCoast for whole genome sequencing. Libraries were prepared with a single library preparation method based on the Illumina Nextera kit and sequenced using the 2× 150-bp paired-end Illumina NextSeq 2000 platform at SeqCoast. We processed raw genomic reads using fastp 0.23.2 (90) and assembled draft genome sequences using SPAdes 3.15.5 (91), using default parameters for both. See **Table S7** in the supplemental material for all genome sequencing statistics.

### Detection, identification, and clustering of *Achromobacter* BGCs

We identified BGCs in *Achromobacter* spp. using antiSMASH 8.0.4 (92) with the following parameters: --cb-general --cb-knownclusters --cb-subclusters. To group the BGCs into GCFs, we used BiG-SCAPE 2.0.3 (93) with the following parameters: -- mibig-version 4.0 --classify category --mix --include-singletons. We generated a heat map in R using complex Heatmap (94). We used clinker (95) to compare the *alc* BGCs from *Achromobacter* spp. to the reference BGC from *B. pertussis* strain Tohama I (https://mibig.secondarymetabolites.org/repository/BGC0002464.3/index.html).

### Generating *pvdA* complement

We complemented the *pvdA* mutant via integration of *pvdA* using the mini-CTX1 plasmid (96). We amplified *pvdA* using primers oRMS125 (5’-ATGC<u>GGATCC</u>AATGGGAAGCCGACTGATTT-3’) and oRMS116 (5’-ATGC<u>GAATTC</u>CATGCAACGAAAACGCCCTG-3’). These primers introduced BamHI and EcoRI restriction overhangs (underlined), respectively, which we used to introduce *pvdA* into mini-CTX-1. We then transformed the resulting plasmid into *Escherichia coli* SM10 for intergeneric conjugation into *P. aeruginosa*. We selected exconjugants using VBBM agar plates containing 75 µg/mL tetracycline. We passaged exconjugants onto LB plates with 25 μg/ml tetracycline. We confirmed complementation by observing exconjugants under blue light using a transilluminator to detect pyoverdine fluorescence and by PCR using primers oRMS117 (5’-CGCCCCTACGAAGGTACCGACA-3’) and oRMS118 (5’-CGACGGCGGAGTTCAGCTGATA-3’), which anneal up-and downstream of the *attB* site, respectively.

### Generating *pchA* deletion mutants

We deleted the *pchA* gene in *P. aeruginosa* strain PAO1 wild type and Δ*pvdA* using plasmid pEXG2-Δ*pchA* that was provided by Dr. Erika Lutter (Oklahoma State University). We performed intergeneric conjugation using *E. coli* SM10 using VBBM agar plates containing 60 µg/ml gentamicin. We then passaged exconjugants on LB with 15% (wt/vol) sucrose for counterselection. We confirmed deletion of *pchA* using primers *pchA* seq F (5’-CAGCGAGCTACTCGAGGACG-3’) and *pchA* seq R (5’-GTCATGGCCCGTGGGGTTAA-3’).

### MALDI-QqTOF-MSI

We cocultured *A. dolens* and three spots *P. aeruginosa* on 10 mL SCFM with 25 *µ*M FeCl_3_ agar plates at 37°C for 24 h. Following incubation, we excised the coculture from the plate using a sterile razor blade and transferred it to a ground stainless steel MALDI target plate, ensuring that no air bubbles formed between the plate and agar. Afterwards we applied a 1:1 mixture of α-cyano-4-hydroxycinnamic acid and 2,5-dihydroxybenzoic acid (CHCA:DHB) MALDI matrix with a 53-micron stainless steel sieve. We dried the plates at 37°C for 4 h using a homemade spinning apparatus (97).

Subsequently, we removed excess MALDI matrix using compressed air. We spotted 1 *µ*L of saturated red phosphorous in 50% (vol/vol) acetonitrile as a calibrant and allowed it to dry. We performed acquisition on a timsTOF flex (Bruker Daltonics) using timsControl v6.0.3_abe6ecc8_1 and flexImaging v7.5 in positive mode at a lateral resolution of 200 µm. We provide the method parameters in **Table S8** in the supplemental material. We analyzed the resulting .slx files using SCiLS Lab v2026b core. We applied total ion count (TIC) normalization. We set the interval width to ± 4.6 ppm for the resulting ion images and chose peak area as the Interval Processing Mode. We generated the ion image using Toggle Image Blend Mode. All analyses were done in biological duplicate.

### MALDI-QqTOF-DDA-MS^2^

Following incubation as above, we spotted a single colony of *A. dolens* or *P. aeruginosa* from each coculture onto a ground stainless steel MALDI target plate using a sterile toothpick. Immediately after drying, we spotted 1 µL of a 1:1 mixture of CHCA:DHB MALDI matrix with 0.1% (vol/vol) trifluoroacetic acid on top of each colony. We performed acquisition on the same instrument as above. After performing a first scan per each culture spot, we used the resulting .run file to generate a precursor list using flex_MSMS_AutoXecute_Generator.exe (98). We then used the resulting list of 100 precursors, not including the media or the matrix, to schedule DDA at the MS^2^ level. We provide the method parameters in **Table S9** in the supplemental material. We converted the resulting .d files to .mzML files using MSConvert (99) with PeakPicking on the MS level = 1 – 2 and uploaded to GNPS2 (100) for generation of molecular networks. We merged single MS^2^ spectra and converted them to .mgf files using MZmine 3 v4.8.30 (101) and analyzed the resulting files with SIRIUS 4 v6.3.3 (102) for putative annotation of observed signals. All analyses were done using three individual colonies as biological replicates.

### Transcriptomic comparison

We cultured *P. aeruginosa* strain PAO1 monocultures and cocultures with *A. dolens* strain RMS0405 on SCFM + 25 *µ*M FeCl_3_, as above. After 12 h coculture, we used a sterile 1 *µ*L inoculation loop to collect *P. aeruginosa* biomass next to *A. dolens* or an equivalent amount of biomass from monoculture control plates. We resuspended the biomass in 100 *µ*L TE buffer containing 50 mg/mL lysozyme and incubated the sample at 37°C for 30 min, with vortexing at 10 min intervals. We then added 150 *µ*L of Monarch StabiLyse DNA/RNA buffer (NEB) and mixed the samples vigorously. We centrifuged the samples at 16,000 × g for 2 min at ambient temperature. We then extracted RNA from the supernatant using a Monarch Spin RNA Isolation Kit (NEB), following the manufacturer’s instructions. We submitted RNA samples to the Oklahoma State University Microbiomics and Culturomics Core Facility (MCCF). RNA integrity and quality were determined using an Agilent RNA ScreenTape system on a TapeStation 4200, and RNA concentration was determined using the Qubit RNA Broad Range Assay Kit (Invitrogen). RNA-seq libraries were then prepared using the Zymo-Seq RiboFree Total RNA Library Kit (Zymo Research). The samples were then sequenced on an Illumina NextSeq 2000 platform using a NextSeq 2000 P3 100-cycle kit in a paired-end 2 × 50 bp configuration in the Oklahoma State University Next Generation Sequencing Core Facility. A library pool at a concentration of 740 pM was loaded for sequencing, with 5% PhiX included as a positive control.

We trimmed reads with fastp 0.23.2 and used kallisto 0.52.0 (103) to pseudoalign reads against a *k* = 31 index built from *P. aeruginosa* strain PAO1 coding sequencing using 100 bootstrap replicates. We used tximport 1.34.0 to import the abundances for testing differential expression with DESeq2 1.46.0 (104). We retained all genes with a pseudocount ≥10. We used apeglm 1.28.0 (105) to shrink the log_2_ fold changes for ranking and display. We assessed significance using a Wald test on the original log_2_ fold changes using a BH-adjusted *P*-value threshold of 0.05 for significance. To identify potential FBS sequences within promoters, we built a position weight matrix with MEME (106) in MEME suite 5.5.9 from 26 experimentally confirmed *P. aeruginosa* FBS. We then scanned from 300 bp upstream of start codons down to 50 bp within coding sequences for the presence of this sequence motif using fimo with a *P*-value threshold of 10^-4^. We compared the proportion of genes with FBS enriched in each culture condition using a jackknife approach where we left one sample out of each analysis. We tested gene set enrichment using a combination of PseudoCAP categories along with some manually curated gene sets using a one-sided Fisher’s exact test with Haldane-Anscombe-corrected OR and BH-adjusted *P*-values.

### Proteome comparison

We used a protocol we previously employed to extract proteins embedded in agar (107). Briefly, we cultured *P. aeruginosa* strain PAO1 monocultures and cocultures with *A. dolens* strain RMS0405 on SCFM + 25 *µ*M FeCl_3_, as above. After 24 h coculture, we used clean razor blades to excise the agar ∼0.1 cm from the *P. aeruginosa* streaks, without disturbing the bacterial growth. We centrifuged the agar slices at 4,000 × g for 10 min at ambient temperature and then combined them with an equal volume of extraction buffer (50 mM Tric-HCl, 0.2% [wt/vol], 0.2% [wt/vol] *N-*lauroylsarcosine sodium salt, pH 7.5), and incubated the mixture on a shaking platform at 150 RPM overnight. We then centrifuged the samples at 4,000 × g for 20 min at ambient temperature and filtered the supernatant through cheesecloth to remove agar. We added trichloroacetic acid to a final concentration of 7.5% (wt/vol), precipitated protein at −20 °C overnight, and pelleted the precipirated protein by centrifuging the samples at 4,000 × g for 20 min at 4 °C.

The protein pellets were washed three times with cold acetone and resuspended in 150 *µ*L of 5% (wt/vol) sodium dodecyl sulfate (SDS) in 100 mM Triethylammonium bicarbonate (TEAB) (pH 8). The samples were sonicated for 10 min in a refrigerated ultrasonic bath, centrifuged at 16,000 × g for 10 min at 4 °C, and the supernatant was transferred to a clean tube. After quantification using a bicinchoninic acid assay, 50 µg of protein was then transferred to a clean tube and reduced with 10 mM tris(2-carboxyethyl)phosphine at for 15 min at 55 °C. The reduced proteins were then alkylated with 40 mM iodoacetamide in the dark for 10 min and then acidified by adding 11 *µ*L of 27.5% (wt/vol) phosphoric acid. The proteins were precipitated after addition of 913 *µ*L S-Trap binding buffer (90% [vol/vol] methanol with 100 mM TEAB, pH 8) and loaded into an S-Trap Spin column. Then, 40 *µ*L of 50 mM TEAB (pH 8) containing 1 µg MS-grade trypsin/LysC was added and incubated overnight at 37 °C . The peptides were eluted following the manufacturer’s protocol, dried *in vacuo*, and reconstituted in 200 µL using 0.1% (vol/vol) trifluoroacetic acid (TFA).

Subsequently, the peptides were desalted using a C18 peptide desalting spin column equilibrated with a 3% (vol/vol) acetonitrile (ACN) and 0.1% (vol/vol) TFA solution. The bound peptides were washed with the same solution to remove salts and other hydrophilic contaminants before being eluted using a 60% (vol/vol) ACN and 0.1% (vol/vol) TFA solution. The eluted proteins were dried *in vacuo* and reconstituted in 0.1% (vol/vol) formic acid. The peptides were injected onto a 75 µm × 50 cm nano-HPLC column packed with 1.9 µm C18 particles and connected to an Easy-nLC 1200 nano-HPLC system. Peptides were separated by gradient chromatography using 0.1% aqueous formic acid as mobile phase A and 80:20:0.1 ACN/water/formic acid as mobile phase B. Peptide separation was performed using a gradient of 4–32% mobile phase B over 120 min at a flow rate of 250 nL per min. The eluted peptides were ionized using a Nanospray Flex ion source and analyzed on an Orbitrap Fusion Tribrid mass spectrometer (Thermo Fisher Scientific). Mass spectrometric data were acquired in DDA mode using a top-speed scan cycle. Full MS scans were acquired in the Orbitrap, followed by quadrupole selection of precursor ions, higher-energy collisional dissociation (HCD), and MS^2^ analysis of fragment ions in the ion trap.

Raw mass spectrometry files were processed using MaxQuant v2.0.2.0 (108) and searched against the *P. aeruginosa* proteome from the UniProt (accession: UP000002438). The Database searching was performed using standard MaxQuant parameters, with trypsin/P specified as the proteolytic enzyme and up to two missed cleavages allowed. Peptide-spectrum matches and protein identifications were filtered at a false discovery rate (FDR) of 1%. In total, we analyzed the proteomes from two biological replicates with three technical replicates.

We filtered the protein groups from MaxQuant to remove any matches to a decoy database, potential contaminants, and proteins identified with only a modified peptide. We only retained proteins with at least one non-zero label-free quantification (LFQ) intensity and at least two MS^2^ counts. We replaced zero LFQ values with the 1st percentile of the measured LFQ value within a sample. All samples within a biological replicate (three cocultures and three monocultures) were processed together for proteomics, but the two biological replicates were processed and analyzed by mass spectrometry in two separate runs. We accounted for batch effects by fitting each protein with its log_2_ LFQ ∼ intercept + condition + batch. We tested condition coefficients using a two-sided *t*-test on the residual degrees of freedom. We adjusted *P*-values using the BH procedure. We classified proteins as extracellular based on curation of the subcellular localization assignments from the *Pseudomonas* Genome Database (109).

### Statistical analysis and data visualization

We performed all statistical analyses in R with specific packages as described in the preceding sections. We generated graphics using ggplot2 (110) with some cleanup and figure assembly in InkScape.

## Data availability

The raw reads and genome sequences of *Achromobacter* spp. generated during this study have been deposited under BioProject Accession number PRJNA1544629. All MALDI-QqTOF-MSI datasets are deposited at MassIVE under accession MSV000102382. All transcriptomic raw reads are deposited under BioProject Accession number PRJNA1545112. All other raw data and derived data are available at https://doi.org/10.6084/m9.figshare.34702161. The scripts necessary to replicate this work are available at https://github.com/reedstubbendieck/Achromobacter_Pseudomonas_Interactions.

## Acknowledgements

We thank Sandra Armstrong (University of Minnesota) for generous insight into alcaligin. We thank Matthew Cabeen (Oklahoma State University [OSU]), Ryan Hunter (State University of New York at Buffalo), Erika Lutter (OSU), and Avishek Mitra (OSU) for strains. We thank Yingxin Zhang (OSU) for assistance with deletion of *pchA*. We thank Mostafa Elshahed and Aymen Yassir (OSU Microbiomics and Culturomics Core Facility) for assistance with transcriptomics (P20 GM152333). We thank Uma Aryal (OSU) and Ali Ali (OSU) for assistance with proteomics.

This research was supported by the National Institute of General Medical Sciences Centers of Biomedical Research Excellence program (P20 GM152333) to R.M.S. L.R.V. was supported by the 2024 Activating Innovative Graduate Research Award (AIGRA) at the University of California, Santa Cruz. The funders had no role in study design, data collection and interpretation, or the decision to submit the work for publication.

## Author Contributions

**Conceptualization**: Niladri Bhowmik and Reed M. Stubbendieck

**Data Curation**: Niladri Bhowmik, Laura Rodriguez-Velandia, Laura M. Sanchez, Reed M. Stubbendieck

**Formal analysis**: Niladri Bhowmik, Laura Rodriguez-Velandia, Laura M. Sanchez, and Reed M. Stubbendieck

**Funding acquisition**: Laura Rodriguez-Velandia and Reed M. Stubbendieck

**Investigation**: Niladri Bhowmik, Laura Rodriguez-Velandia, Mercedes Pérez Pérez, Sydney M. Morabbi, Laura M. Sanchez, and Reed M. Stubbendieck

**Resources**: Reed M. Stubbendieck

**Supervision**: Laura M. Sanchez and Reed M. Stubbendieck

**Visualization**: Niladri Bhowmik, Laura Rodriguez-Velandia, and Reed M. Stubbendieck

**Writing – original draft**: Niladri Bhowmik and Reed M. Stubbendieck

## Supplemental Material

**Figure S1. The severity of *Achromobacter*-*Pseudomonas* coinfections increases with pathogen relative abundance.** (**A**) Effect of coinfection of *Pseudomonas* and *Achromobacter* on FEV_1_ pp after adjustment for age and BioProject, repeated across different relative abundance thresholds. Samples missing either of these values were excluded. Points and error bars show the average change in adjusted FEV_1_ pp and 95% CI. Each estimate is the change relative to *Pseudomonas*-positive samples lacking *Achromobacter*, with the other pathogens held constant. The dashed vertical line marks no difference. BH-corrected *P*-values from per-coefficient *t*-tests are indicated to the right of each comparison. (**B**) Among pwCF coinfected with *Achromobacter* and *Pseudomonas*, lung function versus combined relative abundance of *Achromobacter* or *Pseudomonas*. The pies represent the relative abundance of *Achromobacter* (dark-red), *Pseudomonas* (yellow-green), and other bacteria (gray). Pies are jittered to avoid overplotting. The dashed vertical line separates samples that are pathogen-dominated on the right from those that are not on the left. The line is the ordinary least squares fit with its 95% confidence band. Spearman’s ρ and *P*-value are indicated at top-right.

**Figure S2. The *alc* BGC of *Achromobacter* spp. are highly conserved**. Comparison of *alc* BGCs from *Achromobacter* spp. and *B. pertussis* strain Tohama I (BGC0002464). Darker shading corresponds to higher sequence identity, as indicated by the scale below. Each color corresponds to a distinct homologue. The genes in the *alc* BGC are labeled. Scale bar is 1.0 kb.

**Figure S3. Alcaligin biosynthesis genes are conserved in *Achromobacter* spp.** (**A**) Biosynthetic pathway for alcaligin from L-ornithine. (**B**) Heatmap showing sequence identity of genes required to synthesize alcaligin across *Achromobacter* spp. compared to *B. pertussis* strain Tohama I. A gene was considered present when its best hit spanned at least 70% of the reference length at ≥30% identity, and genes failing these thresholds are drawn as white tiles marked with an X.

**Figure S4.** Mass spectrometry imaging of alcaligin analogues produced during coculture of *A. dolens* **and *P. aeruginosa*.** Mass spectrometry imaging of coculture of *A. dolens* strain RMS0405 (*Ad*, right) and *P. aeruginosa* strain PAO1 on SCFM + 25 *µ*M FeCl_3_. (**A-B**) False-colored blue light image of *P. aeruginosa* (**A**) wild type and (**B**) Δ*pvdA* and *A. dolens*. (**C-D**) Extracted ion images of [M+H]^+^ for alcaligin in coculture between *P. aeruginosa* (**C**) wild type and (**D**) Δ*pvdA* and *A. dolens*. (**E-F**) Extracted ion images of [M+H]^+^ for molecule that may be alcaligin in coculture between *P. aeruginosa* (**E**) wild type and (**F**) Δ*pvdA* and *A. dolens*. (**G-H**) Extracted ion images of [M+H]^+^ for molecule that may be an alcaligin analogue in coculture between *P. aeruginosa* (**G**) wild type and (**H**) Δ*pvdA* and *A. dolens*. Note, the ion intensity for a given molecular feature is comparable between the cocultures with *P. aeruginosa* wild type (left images) and Δ*pvdA* (right images), but not between different ions due to differences in ionization efficiency. All *m/z* values are ± 4.6 parts per million (ppm). Scale bar is 5 mm. Experiment is representative of duplicate samples.

**Figure S5.** Double pyochelin and pyoverdine biosynthesis mutant exhibits a minor growth defect with cultured with *A. dolens* strain RMS0405. (A) Coculture of *A. dolens* strain RMS0405 (right) and *P. aeruginosa* strain PAO1 (left) on SCFM with no supplemented FeCl_3_. (B) Plate from panel A placed in a blue light transilluminator. (C) Cultures of *P. aeruginosa* growing on SCFM with no supplemented FeCl_3_. (D) Plate from panel **C** placed in a blue light transilluminator. Strains are labeled on the left: wild type (RMS290), Δ*pvdA* (RMS0289), Δ*pchA* (RMS0494), and Δ*pvdA* Δ*pchA* (RMS0489). All images are representative of duplicate samples taken under a blue light transilluminator. Scale bar is 5 mm.

**Figure S6. Mass spectrometry imaging of known *P. aeruginosa* secondary metabolites produced during coculture with *A. dolens*.** Mass spectrometry imaging of coculture of *A. dolens* strain RMS0405 (Ad, right) and *P. aeruginosa* strain PAO1 on SCFM + 25 *µ*M FeCl_3_. **(A-B)** False-colored blue light image of *P. aeruginosa* **(A)** wild type and **(B)** Δ*pvdA* and *A. dolens*. **(C-D)** Extracted ion images of molecular features ([M+H]^+^) for pyoverdine in *P. aeruginosa* **(C)** wild type and **(D)** Δ*pvdA*. **(E-F)** Extracted ion images of [M+H]^+^ for pyochelin in coculture between *P. aeruginosa* **(E)** wild type and **(F)** Δ*pvdA* and *A. dolens*. **(G-H)** Extracted ion images of [M+H]^+^ for pyocyanin in coculture between *P. aeruginosa* **(G)** wild type **(H)** and Δ*pvdA* and *A. dolens*. **(I-J)** Extracted ion images of [M+H]^+^ for phenazine-1-carboxylic acid in coculture between *P. aeruginosa* **(I)** wild type and **(J)** Δ*pvdA* and *A. dolens*. **(K-L)** Extracted ion images of [M+H]^+^ for phenazine dicarboxylic acid in coculture between *P. aeruginosa* **(K)** wild type and **(L)** Δ*pvdA* and *A. dolens*. **(M-N)** Extracted ion images of [M+H]^+^ for phenazine-1-carboxamide in coculture between *P. aeruginosa* **(M)** wild type and **(N)** Δ*pvdA* and *A. dolens*. **(O-P)** Extracted ion images of [M+H]^+^ for Pseudomonas quinolone signal in coculture between *P. aeruginosa* **(O)** wild type and **(P)** Δ*pvdA* and *A. dolens*. **(Q-R)** Extracted ion images of [M+H]^+^ for 2-heptyl-4-quinolone in coculture between *P. aeruginosa* **(Q)** wild type and **(R)** Δ*pvdA* and *A. dolens*. **(S-T)** Extracted ion images of [M+H]^+^ for Rha-Rha-C10-C10 in coculture between *P. aeruginosa* **(S)** wild type and **(T)** Δ*pvdA* and *A. dolens*. Brighter pixels indicate higher signal of the corresponding [M+H]^+^. Note, the ion intensity for a given molecular feature is comparable between the cocultures with *P. aeruginosa* wild type (left images) and Δ*pvdA* (right images), but not between different ions due to differences in ionization efficiency. All m/z values are ± 4.6 parts per million (ppm). Scale bar is 5 mm. Experiment is representative of duplicate samples.

**Figure S7. Assignment of molecular features produced by *A. dolens* strain RMS0405 and *P. aeruginosa* strain PAO1 as siderophores.** Evidence for the assignment of (**A**) *m/z* 1333.6177 as pyoverdine E, (**B**) *m/z* 405.1866 as alcaligin, and (**C**) *m/z* 325.0692 as pyochelin.

**Table S1.** Accessions and metadata for amplicon sequencing samples analyzed in this study.

**Table S2.** Individual growth and pyoverdine production results from cocultures between *A. dolens* and 48 clinical *Pseudomonas* isolates used in this study.

**Table S3.** Expression data for all genes in *P. aeruginosa* strain PAO1 in monoculture and during coculture with *A. dolens* strain RMS0405.

**Table S4.** Predicted genes in *P. aeruginosa* strain PAO1 with FBS-containing promoters.

**Table S5.** Differentially expressed genes in *P. aeruginosa* strain PAO1 related to iron in monoculture and during coculture with *A. dolens* strain RMS0405.

**Table S6.** Proteins detected in *P. aeruginosa* strain PAO1 during monoculture and during coculture with *A. dolens* strain RMS0405. The localization of each protein is indicated in the table.

**Table S7.** Metadata for *Achromobacter* spp. used in this study. Genomic sequencing statistics are provided for strains sequenced as part of this study.

**Table S8.** MALDI-TOF-MSI method parameters for acquisition.

**Table S9.** MALDI-DDA-MS method parameters for acquisition.

